# DDX3X and DDX3Y constitutively form nano-sized RNA-protein clusters that foster enzymatic activity

**DOI:** 10.1101/2023.11.29.569239

**Authors:** Amber Yanas, Him Shweta, Michael C. Owens, Kathy Fange Liu, Yale E. Goldman

## Abstract

DEAD-box helicases, which are crucial for many aspects of RNA metabolism, often contain intrinsically disordered regions (IDRs), whose functions remain unclear. Using multiparameter confocal microscopy, we reveal that sex chromosome-encoded homologous RNA helicases, DDX3X and DDX3Y, form nano-sized RNA-protein clusters (RPCs) that foster their catalytic activities *in vitro* and in cells. The IDRs are critical for the formation of these RPCs. A thorough analysis of the catalytic cycle of DDX3X and DDX3Y by ensemble biochemistry and single molecule photon bursts in the confocal microscope showed that RNA release is a major step that differentiates the unwinding activities of DDX3X and DDX3Y. Our findings provide new insights that the nano-sized helicase RPCs may be the normal state of these helicases under non-stressed conditions that promote their RNA unwinding and act as nucleation points for liquid-liquid phase separation under stress. This mechanism may apply broadly among other members of the DEAD-box helicase family.

## Introduction

DEAD-box family RNA helicases are key regulators of RNA metabolism (1). All DEAD-box helicases contain two highly conserved RecA-like helicase domains flanked by variable N- and C-terminal sequences (2, 3). These flanking regions are often disordered and are thought to both contribute to catalytic activity and facilitate protein-protein and protein-RNA interactions. However, how these terminal sequences affect DEAD-box activity and organization has yet to be determined for many family members. We have selected DDX3X and its sexually dimorphic homolog, DDX3Y (4, 5), to understand further how the flanking regions affect protein function. Although these proteins have ∼92% identical overall amino acid sequences, differences are enriched in their N-terminal intrinsically disordered regions (IDRs). Despite this high level of sequence identity, DDX3X has higher enzymatic activity than DDX3Y (6). With these characteristics, the two homologs provide an ideal system to study how IDRs influence the activities of DEAD-box helicases.

We employed a novel multiparameter confocal spectroscopy-based approach to study these aspects in detail. By mixing DDX3X or DDX3Y with fluorescently labeled double-stranded RNA (dsRNA), this technique enabled us to record single molecule fluorescence bursts to obtain a variety of signals simultaneously, including fluorophore stoichiometry, fluorescence resonance energy transfer (FRET), fluorescence lifetime, time-resolved fluorescence anisotropy decay, and dual color Fluorescence Cross-Correlation Spectroscopy (dcFCCS). dcFCCS provides quantitative information on molecular interactions to determine their kinetics and stoichiometry, diffusion rates, the concentrations of fluorescent molecules, and their hydrodynamic radii beyond the optical diffraction limit (7, 8). With this technique, we comprehensively investigated multiple steps along the full-length DDX3X and DDX3Y catalytic cycle and several IDR truncation variants.

Surprisingly, our results showed that, in the presence of RNA, DDX3X and DDX3Y formed large assemblies at concentrations as low as 10 nM (far below the threshold of liquid-liquid phase separation, LLPS). These clusters exist both *in vitro* and within cells, and their formation is driven by their IDRs, as the removal of both IDRs suppressed cluster formation. IDR truncation also decreased the RNA unwinding and ATPase activity of DDX3X and DDX3Y, suggesting a strong correlation between cluster formation and catalytic activity. Furthermore, we show that the N-terminal IDRs were largely responsible for differentiating the enzymatic activities of DDX3X and DDX3Y but were equally crucial for RPC formation, while the C-terminal IDRs were equivalently important for the enzymatic activities of both proteins but differentiated their cluster formation. Additionally, the results suggest that the faster kinetics of DDX3X relative to DDX3Y is likely due to quicker strand release after unwinding, a main reaction step limiting catalytic turnover. Together, this study advances our understanding of how DDX3X and DDX3Y differ in their functions and provides new insights into how N- and C-terminal flanking regions contribute to the catalytic activity of DEAD-box family helicases.

## Results

### DDX3X and DDX3Y form nano-sized RNA-protein clusters even at nanomolar concentrations

To investigate the assembly state of DDX3X and DDX3Y while performing RNA unwinding, we used dcFCCS, illustrated in **Figure 1A**, to measure the diffusion constant and apparent hydrodynamic radius of actively functioning RNA-protein complexes. It was previously shown via sedimentation analysis and confocal microscopy that at threshold concentrations of ∼5 and ∼3 μM, respectively, DDX3X and DDX3Y phase separate into liquid-like droplets (6). However, the assembly state of DDX3X and DDX3Y at lower concentrations was unknown. We employed a previously described fluorescent-labeled dsRNA target construct (6, 9), having an 18 base pair (bp) double-stranded segment and a 42 nucleotide (nt) 5’ single-stranded poly-U overhang (termed **dsRNA I** here) (**Figure 1A**). The longer overhanging strand was labeled with 5’ Cy3 and the shorter hybridized strand contained 3’ Alexa647 (A647). Fluorescent molecules diffused freely in and out of the fixed-position ∼1 fL detection volume of a confocal microscope with Pulse-Interleaved Excitation (PIE) alternating λ = 532 and 640 nm laser wavelengths, producing a burst of fluorescent photons as each single fluorescent complex traversed the beam (**Figure 1B and 1C**). The occupancy time, *τ*_D_, in the beam, is related inversely to the diffusivity according to *D* = *w_0_ ^2^*/4*τ*, where *w_0_* is the detection volume radius at the beam waist. By cross-correlating the photon bursts of the directly excited Cy3 and A647 emission streams using FRETBursts software (10) from 2.5 to 5-minute recording periods, we quantified the diffusion constant of particles containing both probes.

**Figure 1.**
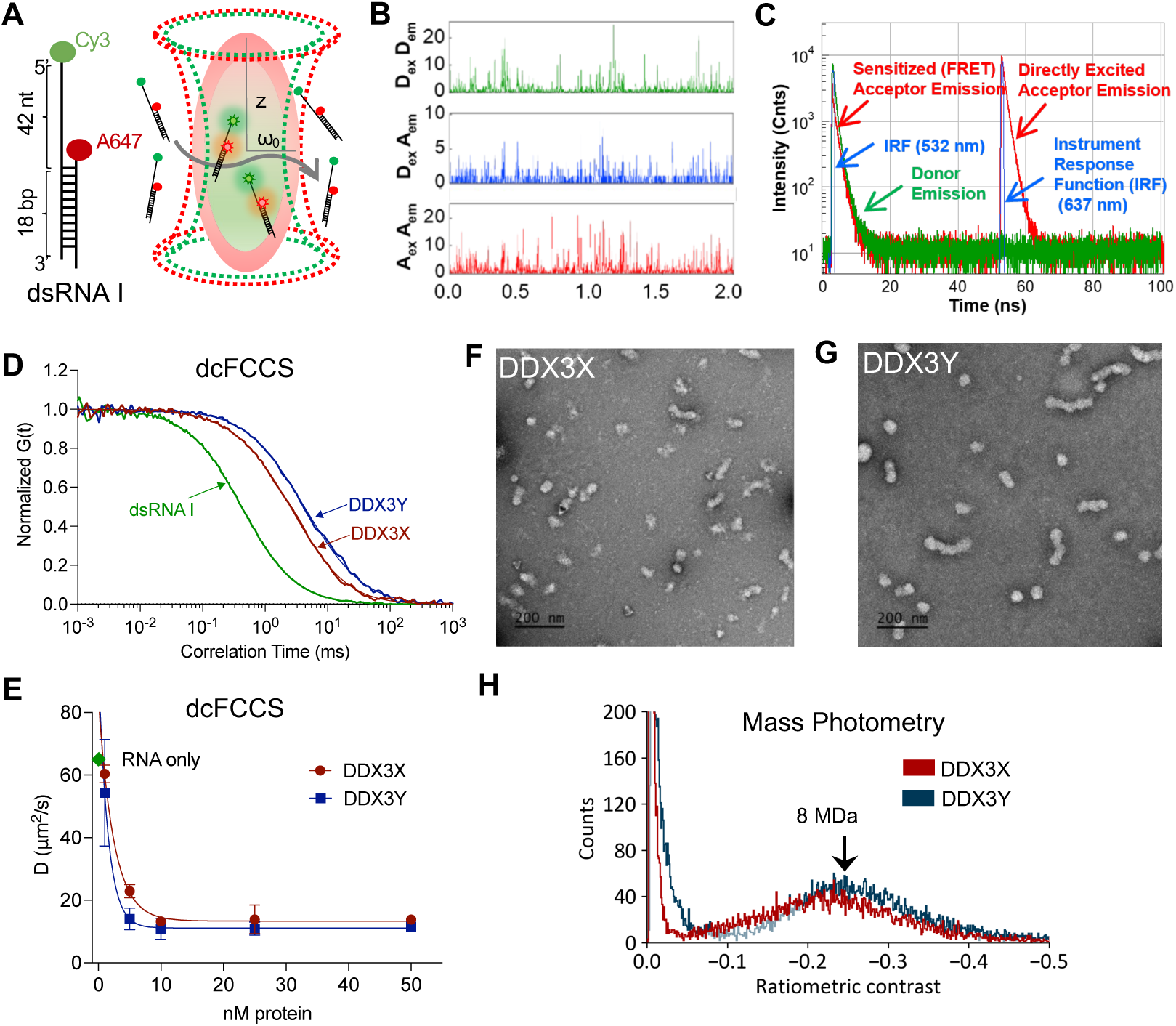
DDX3X and DDX3Y form nano-sized RNA-protein clusters *in vitro*. (A) Schematic of the fluorescent dsRNA I (left) and the confocal detection volume (right). (B) Representative intensity time traces of donor emission upon donor excitation (D_ex_D_em_, green), acceptor emission upon donor excitation (D_ex_A_em_, blue), and acceptor emission upon acceptor excitation (A_ex_A_em_, red). (C) Representative time-correlated single photon counting histograms of the photon distributions over time for D_ex_D_em_ (donor emission, green), D_ex_A_em_ (sensitized acceptor emission, red), and A_ex_A_em_ (directly excited acceptor emission, red). (D) Normalized cross-correlation curves of fluorescent dsRNA I alone (green), full-length MBP-DDX3X (red) and MBP-DDX3Y (blue). (E) Diffusion coefficients (µm^2^/s) of increasing nM concentrations of MBP-DDX3X and DDX3Y with the fluorescent RNA reporter (dsRNA I). Values were fit to an exponential decay equation using GraphPad Prism 9. Curves represent three independent measurements for each point, error bars represent SEM. (F) Representative electron microscopy image of negatively stained 1 μM full-length MBP-DDX3X with 1 nM dsRNA I. Scale bar, 200 nm. (G) Representative electron microscopy image of negatively stained 1 μM full-length MBP-DDX3Y with 1 nM dsRNA I. Scale bar, 200 nm. (H) Histograms of ratiometric contrast of 1 µM of full-length MBP-DDX3X and MBP-DDX3Y were obtained using mass photometry. An average ratiometric contrast of -0.25 corresponds to a molecular weight of ∼8 MDa.

dsRNA I, in the absence of any DDX3 proteins, gave *τ*_D_ = 0.45 ± 0.02 ms (mean ± s.e.m., n = 7), corresponding to *D* = 67.8 µm^2^/s for monomeric 27 kDa dsRNA sample. Unexpectedly, the addition of 2 µM MBP-tagged DDX3X or DDX3Y dramatically slowed the occupancy time to *τ*_D_ = 2.4 ± 0.4 and 4.3 ± 1.1 ms (corresponding to *D* = 13.0 and 7.7 µm^2^/s), respectively (**Figure 1D, S1A**). Given that *D* is proportional to the inverse of the hydrodynamic radius for a compact particle and that mass is proportional to the radius ***cubed***, the 5.4 – 2.8-fold changes in *τ*_D_ shown in **Figure 1D** when the protein was added correspond to clusters of RNA and protein (RNP) with masses 140 – 680-fold higher than the original 27 kDa dsRNA construct. The individual DDX3 protein molecules have a molecular mass (*M*_W_) = ∼120 kDa. Thus, the particles exhibiting these huge decreases in dsRNA diffusivity when MBP-DDX3X or MBP-DDX3Y were added correspond to clusters at *M*_w_ ∼3.5 – 20 MDa containing several protein and RNA molecules. At a density of 1.4 g/cm^3^ (11), such particles correspond to ∼20 nm diameter-sized RNA-protein clusters, which we term “nano-clusters” or “RNA-protein clusters (RPCs).” The shape and compactness affect diffusion, but the huge observed increase in diffusion time when the fluorescent dsRNA was mixed with the DDX3 proteins would not be affected enough by these factors to alter the conclusion that the dsRNA is contained in RPCs with remarkably increased mass. 2 µM MBP protein alone (without the DDX3s) did not show any binding to the dsRNA substrate or slowing of its diffusion (**Figure S1A**). These surprising observations led us to characterize the nano-clusters in more detail and seek other evidence for sub-diffraction limit clustering of RNA and the helicases described below. Surprisingly low concentrations of DDX3X and DDX3Y formed RPCs with the dsRNA I substrate: the values of *τ*_D_ and *D* reached their saturating low values at only 10 nM MBP-DDX3X or MBP-DDX3Y, with MBP-DDX3Y giving a stronger effect (**Figure 1E**). At 1 – 5 nM DDX3 concentrations, the cross-correlation curves showed evidence of mixtures of rapidly diffusing free dsRNA and slower diffusing nano-clusters (**Figure S1B and S1C**).

Full-length DDX3X and DDX3Y purified from bacteria co-purify with cellular RNA (6). Thus, the RPCs described above are likely present in the solutions before adding the fluorescent-labeled RNA and they are made apparent in the dcFCCS spectra by capturing the labeled dsRNA. To further understand the impact of RNA incorporation in the RPCs, single-probe Fluorescence Correlation Spectroscopy (FCS) experiments were performed with mCherry-tagged DDX3X and DDX3Y and with various concentrations of total RNA extracted from HEK293T cells (**Figure S1D, S1E and S1F**). The DDX3X-mCherry and DDX3Y-mCherry constructs co-purify with about 33% less endogenous cellular RNAs than the MBP-tagged constructs (6), so we supplemented with total RNA from HEK293T cells to induce cluster formation at low protein concentrations. When 50 ng/µL of total RNA was added to 80 nM mCherry-tagged DDX3X or DDX3Y, clusters formed and diffused similarly to the MBP-tagged DDX3 RPCs, which consisted of freely diffusing dsRNA and slowly diffusing RPCs (**Figure S1F**). In control experiments adding RNA to the mCherry protein alone, mCherry diffusion was not slowed (**Figure S1G**). These experiments indicate that RPC formation requires an interaction between DDX3 and RNA.

As an orthogonal approach, we performed transmission electron microscopy (TEM) of negatively stained samples of 1 μM MBP-DDX3X or MBP-DDX3Y in the presence of 1 nM dsRNA I. Both DDX3X (**Figure 1F**) and DDX3Y (**Figure 1G**) formed elongated clusters clearly visible by negative staining, further supporting the dcFCCS observations. As a third approach, we used mass photometry to test for RPC formation. In this optical microscopic interferometric scattering technique (iSCAT), slight increases in image contrast are linearly related to the mass of test particles landing on a microscope cover slip (12). The molecular masses are calibrated against samples with known molecular weights, such as BSA, apo-ferritin, and the human 80*S* ribosome (**Figure S1H and S1I**). At 1 nM fluorescent dsRNA I and 1 µM MBP-DDX3X or MBP-DDX3Y, well below concentrations leading to LLPS, the mass distributions indicated 200 – 600 kDa oligomers and 6 – 12 MDa clusters corresponding well with the dcFCCS and EM findings (**Figure 1H**). Three independent approaches suggest that RNA promotes DEAD-box helicases DDX3X and DDX3Y to form RPCs clusters at concentrations below the threshold of liquid-liquid phase separation. A typical RPC is ∼20 nm in diameter and contains an estimated local concentration of ∼9.5 µg/mL cellular RNA and 285 µg/mL protein.

### The intrinsically disordered regions of DDX3X and DDX3Y drive the formation of RPCs

IDRs are known as major contributors to multivalent protein-protein and protein-RNA interactions (13), which led us to study whether they facilitate the formation of RPCs. We generated constructs with both the N- and C-terminal IDRs removed (ΔΔ DDX3X, a.a. 132-607 and ΔΔ DDX3Y, a.a. 130-605) tagged with MBP (**Figure S2A**). The dcFCCS cross-correlation curves for the ΔΔ constructs indicated that the ΔΔ DDX3X clusters diffused 1.53-fold faster than full-length (FL) DDX3X and ΔΔ DDX3Y diffused 2.26-fold faster than FL DDX3Y (**Figure 2A**). These values suggest that the masses of the ΔΔ particles were 3.5-fold and 11.5-fold smaller for DDX3X and DDX3Y, respectively, than the corresponding FL clusters. The size differences between FL and ΔΔ constructs were next studied using both mass photometry (**Figure 2B**) and TEM (**Figure 2C-F**). Both techniques indicated that assemblies formed from the ΔΔ constructs were drastically smaller than those of the full-length proteins. Mass photometry indicated that ΔΔ DDX3 assemblies consist of three or four monomers; no larger particles were detected.

**Figure 2.**
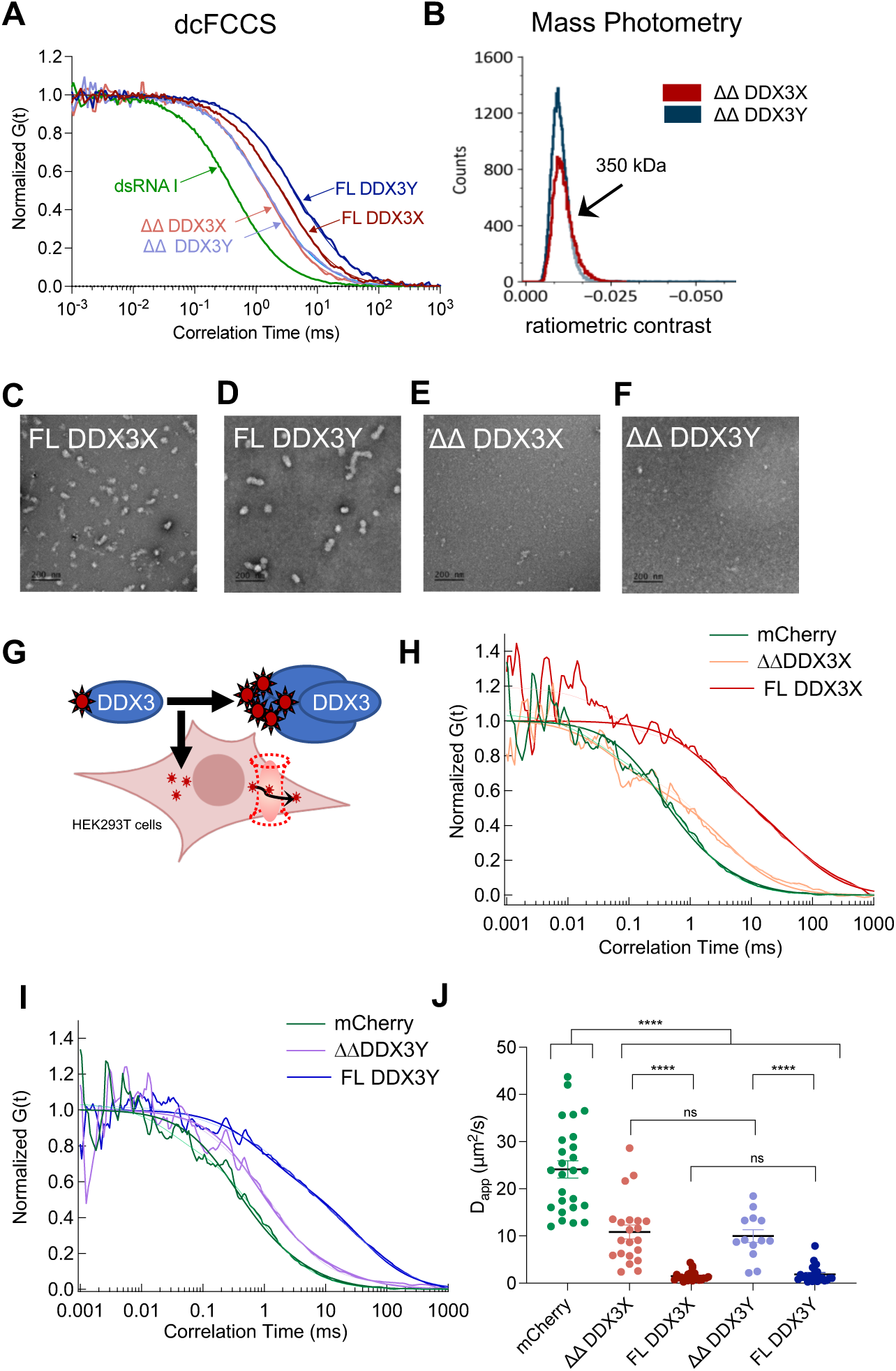
The intrinsically disordered regions of DDX3X and DDX3Y drive the formation of RPCs *in vitro* and in human cells. (A) Normalized cross-correlation curves of fluorescent RNA alone, MBP-tagged ΔΔ DDX3X, ΔΔ DDX3Y, full-length DDX3X and full-length DDX3Y. (B) Histogram of the ratiometric contrasts of ΔΔ DDX3X and ΔΔ DDX3Y obtained through mass photometry, the ratiometric contrast of -0.014 corresponds to a mass of 350 kDa. (C) Representative electron microscopy image of negatively stained 1 μM MBP-tagged FL DDX3X with 1 nM 1 nM RNA I. Scale bar, 200 nm (D) Representative electron microscopy image of negatively stained 1 μM MBP-tagged FL DDX3Y with 1 nM 1 nM RNA I. Scale bar, 200 nm (E) Representative electron microscopy image of negatively stained 1 μM MBP-tagged ΔΔ DDX3X with 1 nM 1 nM RNA I. Scale bar, 200 nm (F) Representative electron microscopy image of negatively stained 1 μM MBP-tagged ΔΔ DDX3Y with 1 nM RNA I. Scale bar, 200 nm (G) Schematic showing the in-cell FCS experiments. (H) Normalized auto-correlation curve of mCherry alone, mCherry tagged ΔΔ DDX3X, and mCherry tagged full-length DDX3X diffusing in live transfected HEK293T cells. (I) Normalized auto-correlation curve of mCherry alone, mCherry tagged ΔΔ DDX3Y, and mCherry tagged full-length DDX3Y diffusing in live, transfected HEK293T cells. (J) Apparent diffusion coefficients (D_app_) (µm^2^/s) of mCherry, ΔΔ DDX3X-mCherry, full-length DDX3X mCherry, ΔΔ DDX3Y-mCherry, full-length DDX3Y-mCherry from live transfected HEK293T cells. Plots and significance were calculated in GraphPad Prism 9 using Student’s two-tailed t-test. ns, p < 0.12, *, p < 0.033, **, p < 0.002. ***, p < 0.0002. ****, p < 0.0001.

### DDX3X and DDX3Y form RPCs in human cells dependent on their IDRs

To assess whether DDX3X and DDX3Y form clusters in cells, we expressed mCherry-tagged FL DDX3X, FL DDX3Y, ΔΔ DDX3X, ΔΔ DDX3Y, and mCherry alone to similar levels in HEK293T cells (**Figure 2G, S2B-C**). These measurements were taken without adding cellular stressors (such as arsenite or sorbitol) that would induce stress granules (6, 14). The diffusion of FL DDX3X-mCherry and FL DDX3Y-mCherry was the slowest, with an average apparent diffusion coefficient *D_app_* = 1.5 ± 0.23 µm^2^/s and 1.8 ± 0.40 µm^2^/s, respectively (mean ± S.E.M.; n = 24 and 26, **Figure 2J**). The truncated mCherry-tagged proteins, with IDRs removed (ΔΔ DDX3X-mCherry and ΔΔ DDX3Y-mCherry), diffused much faster, with *D_app_* = 11 ± 1.5 µm^2^/s and 10 ± 1.4 µm^2^/s, respectively (mean ± s.e.m.; n = 20 and 24, **Figure 2J)**. The diffusion of mCherry alone was the fastest, *D_app_* = 24 ± 1.8 µm^2^/s (mean ± s.e.m.; n = 20 **Figure 2J**). In cell FCS results showed that the diffusion of these proteins in cells (**Figure 2J**) matches their relative diffusion pattern *in vitro* (**Figure 2A**). Collectively, the results support the notion that DDX3X and DDX3Y exist as slowly diffusing RPCs in cells as well as *in vitro*. Given that DDX3X is known to perform many functions in the cell (15–19), these cellular RPCs likely contain a number of other components, such as ribosomes and mRNAs.

### Ensemble measurements reveal specific roles for each IDR in the DDX3 catalytic cycle

To understand how individual IDRs influence catalytic activity, we measured the dsRNA unwinding rates of full-length and truncated proteins via a gel-based unwinding assay (20) using a previously optimized smaller substrate dsRNA: a 13 bp duplex with a 25 nt 3’ overhang (termed **dsRNA II**) (21). When ATP-dependent unwinding activity was assessed with this substrate at 100 nM DDX3X or DDX3Y, DDX3X unwound the substrate faster and to a greater degree than DDX3Y (**Figure 3A-B**). This difference was observed over all measured concentrations of DDX3X and DDX3Y (**Figure 3C**). In this concentration range of DDX3X and DDX3Y, RPCs are observed with the smaller dsRNA II as with dsRNA I (**Figure S3A**). When we removed the IDRs from DDX3X and DDX3Y and repeated the unwinding assay, both ΔΔ truncations had significantly weaker activity than their full-length counterparts. The data suggest that the IDRs promote enzyme activity in addition to promoting the formation of RPCs (**Figure 3D**).

**Figure 3.**
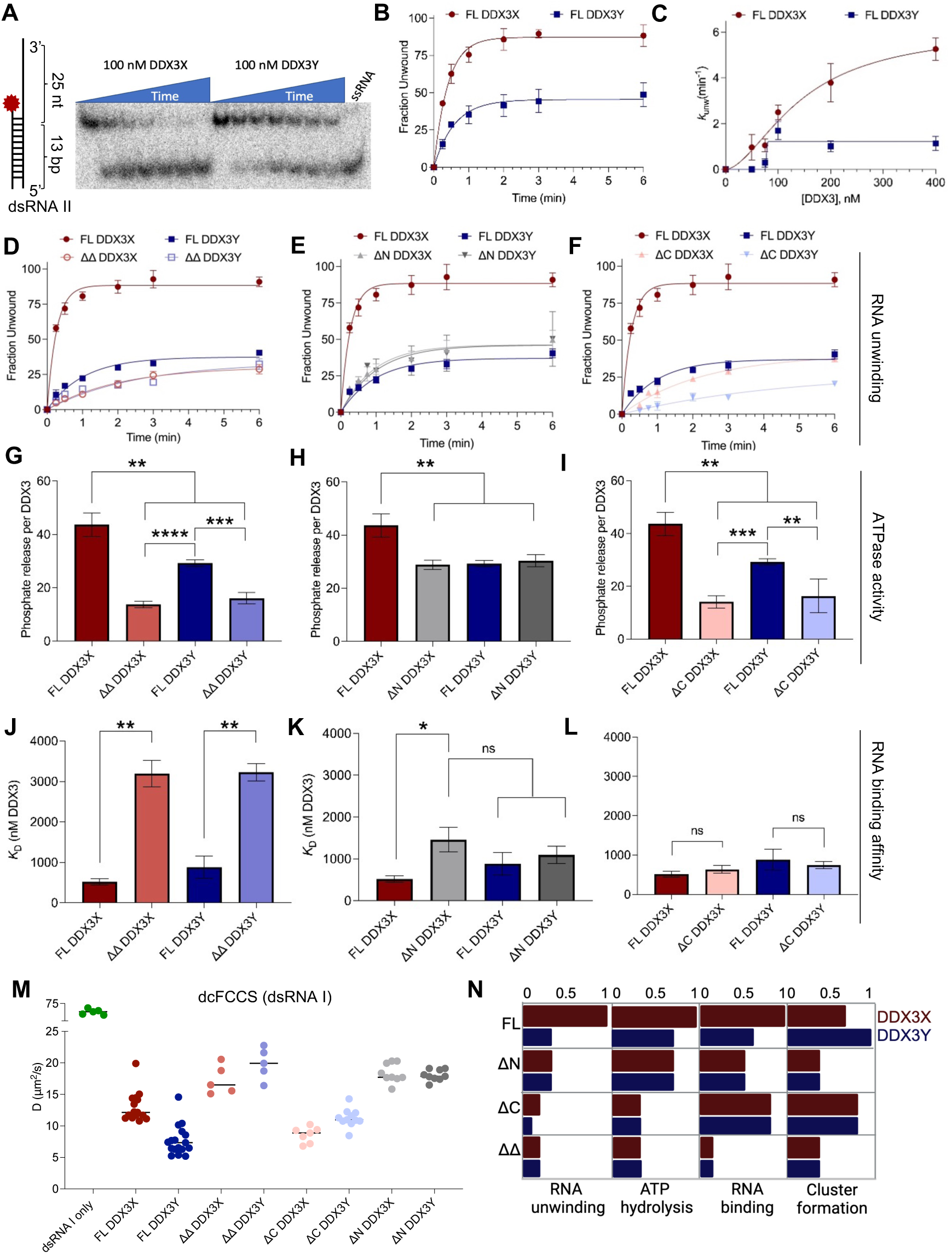
Functional annotation of individual IDRs in forming protein-RNA clusters and promoting enzymatic activities of DDX3X and DDX3Y. (A) Left, schematic of RNA duplex used for gel-based unwinding experiments, contains a 13 nt short strand radiolabeled at the 5’ end annealed to a 38 nt unlabeled long strand. The RNA was annealed to produce a 13 bp duplex with a 25 nt 3’ single-stranded overhand (dsRNA II). Right, representative unwinding gel for 100 nM full-length MBP-DDX3X and MBP-DDX3Y. “ssRNA” sample was generated by taking a final reaction sample and heating it to 95° C for two minutes. (B) Quantification of Fig. 3A. Fraction unwound was quantified and fit as described in the methods. Plots are the average of three independent experiments. (C) Unwinding rates at the indicated concentrations were calculated as in (B) for full-length MBP-DDX3X and MBP-DDX3Y. Values were from three independent experiments and were fit using Hill kinetics. (D-F) Fraction unwound curves comparing full-length MBP-DDX3X/DDX3Y and to the indicated truncation. Values were from three independent experiments. (G-I) Malachite green ATPase assay showing release of phosphate from ATP hydrolysis stimulated by 100 ng/μL HeLa total RNA for full-length MBP-DDX3X/DDX3Y and the indicated truncations. The assay was performed as described in the methods. Values were from three independent experiments. Significance was calculated in GraphPad Prism 9 using the Student’s two-tailed t-test. *, p < 0.05, **, p < 0.01. ***, p < 0.001, ****, p < 0.0001. (J-L) Summarized *K*_D_ values for the indicated constructs, calculated as described in methods. Values were from three independent experiments. Plots and significance were calculated in GraphPad Prism 9 using the Student’s two-tailed t-test. ns, not significant, *, p < 0.05, **, p < 0.01. (M) Diffusion coefficients (µm^2^/s) were obtained by fitting the cross-correlation curves using single component diffusion model for RNA I alone, MBP-tagged full-length DDX3X, full-length DDX3Y, ΔΔ DDX3X, ΔΔ DDX3Y, ΔC DDX3X, ΔC DDX3Y, ΔN DDX3X, and ΔN DDX3Y. These diffusion coefficients correlate with the size of the molecules. (N) Schematic illustrating the role of each IDR in catalysis and cluster formation. Values for FL DDX3X were set to 1 for RNA unwinding, ATP hydrolysis, and RNA binding, and values for other constructs were normalized to those of FL DDX3X. Values for FL DDX3Y from Figure 3M were set to 1 for cluster formation, and values for other constructs were normalized to those of FL DDX3Y. For RNA binding and cluster formation, values were inverted before normalization.

To assess how each IDR contributes to the differences we observe between DDX3X and DDX3Y, we conducted the unwinding assays using constructs lacking either the N- or C-terminal IDR (ΔN and ΔC) (**Figure S2B and S3C**). Removing the N-terminal IDR of DDX3X decreased its dsRNA unwinding rate (**Figure 3E and S3C**) to be equal to that of DDX3Y, while removing the N-terminus of DDX3Y did not alter its unwinding rate at all. Removing the C-terminal IDRs from both proteins caused a sharp decrease in dsRNA unwinding (**Figure 3F and S3C**) down to the same level as the ΔΔ truncations (**Figure 3D**).

Next, we assessed the ATPase activity of each construct by measuring phosphate release from ATP hydrolysis with a malachite green colorimetric assay in the presence of HEK293T total RNA to stimulate activity as previously described (6). The trends for ATPase activity among the truncated constructs mirrored their trends in RNA unwinding rate: removing both IDRs decreased the ATPase activity to the same low level (**Figure 3G**), whereas removing the N-terminal IDRs only dropped the ATPase activity of DDX3X but not DDX3Y (**Figure 3H**). Removing the C-terminal IDRs decreased ATPase activity for both proteins to the level of the double truncation (**Figure 3I**). These results point towards a model in which the N-terminal IDR of DDX3X supports its catalytic activities RNA unwinding and ATPase activities in a manner in which DDX3Y’s N-terminal IDR does not achieve, while the C-terminal IDRs of both proteins are equally supportive of dsRNA unwinding and ATP hydrolysis. These findings align with the facts that the C-terminal IDRs of DDX3X and DDX3Y are similar and most of the sequence differences reside in their N-terminal IDRs, which is sufficient to differentiate fundamental properties of these two helicases.

We assessed the role of each IDR in dsRNA binding via electrophoretic mobility shift assays using dsRNA I (**Figure 3J-L, S3D, S3E**). Full-length DDX3X had a slightly lower *K_D_* value than full-length DDX3Y (519 ± 43 nM vs. 884 ± 155 nM mean ± s.e.m., respectively) (**Figure S3D**). Removing both IDRs (ΔΔ) drastically increased the *K_D_* values more than threefold, in line with the results of the unwinding assays (**Figure 3J**). Removing the N-terminal IDR from DDX3X increased its *K_D_* value significantly while removing the N-terminal IDR from DDX3Y resulted in a much smaller, statistically insignificant, increase (**Figure 3K**). These results mimic the unwinding assay above, where the N-terminal IDR of DDX3X contributed to its overall activity while the N-terminal IDR of DDX3Y did not. In contrast, removing the C-terminal IDRs of either protein resulted in small, statistically insignificant changes in binding affinity, indicating the C-terminal IDRs of both proteins may be dispensable for dsRNA binding in the presence of the N-terminal IDR (**Figure 3L**).

By dcFCCS, it was determined that single N-terminal IDR deletion constructs showed faster diffusion similar to ΔΔ, showing that the N-terminal IDR drives cluster formation (**Figures 3M and S3F**). The single C-terminal IDR deletion constructs showed an intermediate diffusion between FL DDX3X and DDX3Y (**Figure 3M**), showing that the C terminal IDRs of both proteins contribute to their difference in cluster formation. Collectively, these results indicate the N-terminal IDRs of DDX3X and DDX3Y are both major drivers of RPC formation and major differentiators of their enzymatic activities. Conversely, their C-terminal IDRs are equally critical for enzymatic activity but are a differentiator of RPC size for DDX3X and DDX3Y (**Figure 3N**).

### RNA release is a key step endowing the stronger helicase activity of DDX3X relative to DDX3Y

The catalytic activity of DDX3 is critical for controlling gene expression and deficits are linked to many human diseases, including cancer (19, 22, 23). Since DDX3X is a more active RNA helicase than DDX3Y (**Figure 3**), we sought to reveal the kinetic steps determining this difference. The major catalytic steps in the helicase cycle are 1) RNA binding, 2) strand separation, 3) ATP hydrolysis, and 4) RNA release. Additionally, it is possible that DDX3 is capable of some degree of ATP-independent strand separation, as this has been observed for other DEAD-box proteins (24). The differences between DDX3X and DDX3Y at each step can be elucidated by relating single-burst fluorescence spectroscopy and the ensemble biochemical assays.

The interleaved wavelength excitation scheme of the confocal spectrometer enables each burst of photons, corresponding to each diffusing particle, to be categorized according to the relative stoichiometry of the two fluorescent labels on the dsRNA substrate. For particles doubly labeled with donor Cy3 and acceptor A647 (DL), the FRET efficiency can act as a molecular distance ruler for the separation between the two fluorophores ((25) and see Methods). These are displayed as 2D histograms in **Figure 4A-C** with stoichiometry (*S*) in the vertical dimension and FRET efficiency (*E*) in the horizontal. *S* = 1 for Cy3 donor-only (DO, long strand) species, *S* = 0.5 for doubly labeled (DL, duplex) particles with a 1:1 donor:acceptor ratio, and *S* = 0 for particles containing only A647 acceptor (AO, short strand) probes. The 1D histogram to the right of each 2D plot shows the distribution between these three species. The 1D histogram at the top of each panel shows the FRET efficiency distribution for the DL (S ≍ 0.5) group.

**Figure 4.**
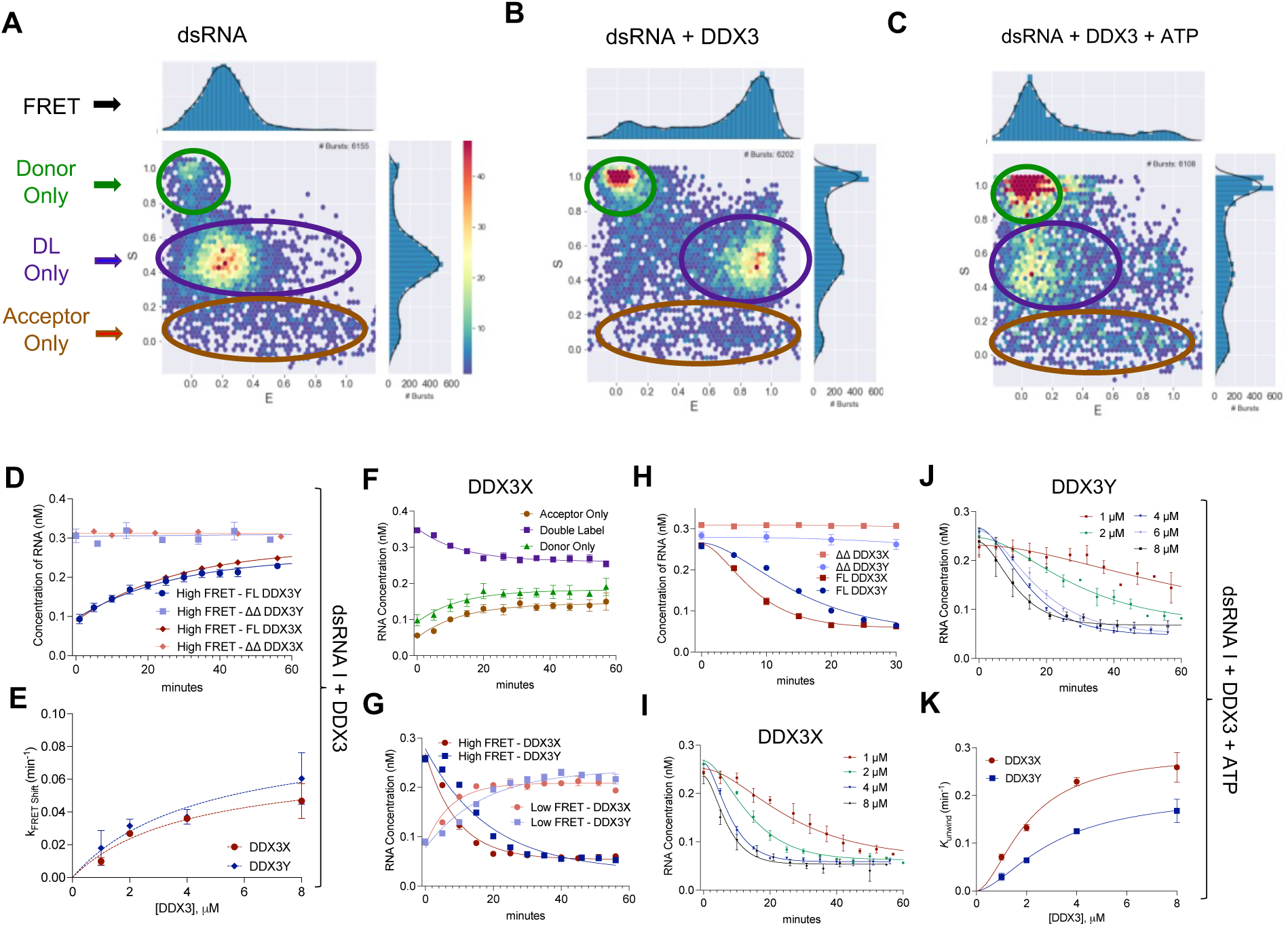
DDX3X has faster unwinding activity compared to DDX3Y, and the IDRs are important for catalytic activity. (A) A 2D plot of *E-S* representation showing DO (Cy3; *S* >0.8), FRET (DL; *S*∼0.5) and AO (Alexa-647; *S*<0.2) populations along with the marginal *E* (X-axis) and *S* (Y-axis) histograms. The left scatter plot (dsRNA only) shows RNA with an abundance of low FRET DL particles and low DO and AO particles before adding the protein. (B) The middle scatter plot shows the particles when DDX3 protein is added, which leads to an abundance of high FRET, and DL particles and some increase in DO and AO particles. (C) The right scatter plot shows the particles when 1 mM ATP is added. There is a shift of the high FRET DL particles to low FRET and an increase in AO and DO particles, indicating unwinding activity. (D) Fraction of DL (Alexa647 and Cy3 double-labeled) particles when 4 µM MBP-tagged full-length DDX3X (red) and full-length DDX3Y (dark blue), ΔΔDDX3X (pink), or ΔΔDDX3Y (light blue) is added to RNA. WT DDX3X and DDX3Y exhibit ATP-independent helicase activity and subsequent reannealing activity. ΔΔDDX3X and ΔΔDDX3Y do not show this activity. Values were fit to an exponential decay equation using GraphPad Prism 9. Curves represent three independent measurements for each point. (E) Concentration-dependent curve of the kinetics of the FRET shift from low to high FRET plotted at various protein concentrations for both MBP-DDX3X (red) and MBP-DDX3Y (blue). Both proteins show similar activity. Values were fit to a Michaelis-Menten equation using GraphPad Prism 9. Curves represent three independent measurements for each point. (F) Example plot showing the RNA concentration of single-stranded acceptor (Alexa647), single-stranded donor (Cy3), and DL (duplex, both Alexa647 and Cy3) with the addition of ATP. Values were obtained by quantifying the respective burst proportion and multiplying by the starting RNA concentration (0.5 nM) see methods. Values were fit to an exponential decay equation using GraphPad Prism 9. Curves represent three independent measurements for each point. (G) Example plot showing the high FRET and low FRET DL populations of both 4 µM full-length MBP-DDX3X (red) and MBP-DDX3Y (blue) multiplied by the starting RNA concentration (0.5 nM). With the addition of ATP, the high FRET population decreases, and the low FRET population increases for DDX3X and DDX3Y. Values were fit to an exponential decay equation using GraphPad Prism 9. Curves represent three independent measurements for each point. (H) Example kinetics curve of 4 µM MBP-tagged full-length DDX3X (red) and DDX3Y (dark blue), ΔΔ DDX3X (pink), ΔΔ DDX3Y (light blue) with 1 mM ATP. RNA concentration (decrease in DL x decreases in high FRET x 0.5 nM RNA, see methods, equations 5 and 6) is plotted on the Y-axis. Values were fit to an exponential decay equation using GraphPad Prism 9. Curves represent three independent measurements for each point. (I) Individual kinetics curves plotted for 1, 2, 4, and 8 µM full-length MBP-DDX3X with 1 mM ATP. RNA concentration is plotted on the Y-axis. Values were fit to an exponential decay equation using GraphPad Prism 9. Curves represent three independent measurements for each point. (J) Individual kinetics curves plotted for 1, 2, 4, and 8 µM full-length MBP-DDX3Y with 1 mM ATP. RNA concentration is plotted on the Y-axis. Values were fit to an exponential decay equation using GraphPad Prism 9. Curves represent three independent measurements for each point. (K) Helicase activity (*K_unwind_*) of DDX3X (red) and DDX3Y (blue) at various protein concentrations. Values were fit to Hill kinetics using GraphPad Prism 9. Curves represent three independent measurements for each point.

DL RNA alone had a moderate *E* = 0.25 FRET efficiency indicating a distance between the two probes averaging approximately 0.63 nm ((6) and **Figure 4A**). When DDX3X or DDX3Y was added, several changes in the RNA sample were observed: 1) the proportion of DL particles decreased slightly, indicating a partial ATP-independent helicase activity (compare the *S*-histograms in **Figure 4A and B**), and then 2) the fraction of DL particles returned partially toward the original value (**Figure 4D**) indicating partial re-annealing of the dsRNA I or re-capture of the labeled strands into the RPCs. 3) FRET efficiency gradually increased to a steady state level of ∼0.9, indicating the two fluorescent probes were much closer with protein bound (**Figure 4B** and (6)). These ATP-independent properties of DDX3X and DDX3Y required the presence of the IDRs (**Figure 4D**). The half saturation value of DL formation and increase in high FRET were quite similar for DDX3X and DDX3Y (*K_M(DDX3X)_* = 4.1 ± 2 µM, *K_M(DDX3Y)_* = 3.7 ± 3 µM) (**Figure 4E, S4A, S4B**).

To directly study the strand annealing activity of the two helicases, we first separately pre-mixed the Cy3-only strand (DO) with protein and the A647-only strand (AO) with protein. The two single strand-DDX3 samples were then mixed together, and the formation of DL particles was measured (**Figure S4C**). The rates and extent of DL formation were quite similar for DDX3X and DDX3Y (**Figure S4C**). Mixing the two single RNA strands without protein or with MBP alone showed little spontaneous annealing at 20 °C (**Figure S4C**). These data reveal that both DDX3X and DDX3Y are capable of a similar degree of ATP-independent RNA unwinding and protein-dependent RNA annealing.

When ATP was added to samples of RNA and DDX3X or DDX3Y (equilibrated after adding the protein to dsRNA as in **Figure 4B**), the proportion of DL particles declined during the next 30 – 60 min indicating separation of the two RNA strands (**Figure 4C, 4F**). As [DL] decreased, [AO] and [DO] increased appropriately (**Figure 4F**). FRET efficiency of the remaining DL population also declined gradually to *E* = 0.1, indicating ATP-dependent strand separation (**Figure 4G**). Note that the low-FRET group that dominates in the DL section of the scatterplot in **Figure 4C** does not represent full strand separation because it is determined solely from the DL particles containing both fluorescent probes. The reduction of *E* in DL clusters is likely due to multiple partial helicase cycles that fail to complete strand separation because of rapid local re-annealing within the cluster. ΔΔ DDX3X and ΔΔ DDX3Y show almost no unwinding activity (**Figure 4H**), supporting the observation by gel analysis that the IDRs are critical for this activity (**Figure 3**). The protein concentration dependence of unwinding rates (**Figure 4I-K**) shows that DDX3X is more effective than DDX3Y in full-strand separation, in line with the results of the gel-based unwinding assay (**Figure 3C**).

Nanosecond time-resolved anisotropy of the fluorescent probes in the free and clustered dsRNA particles provided a method to detect dissociation of the RNA from the protein and RPC clusters (**Figure 5**). Free dsRNA tumbles with a rotational correlation time (*Φ*_r_) of 0.8 ± 0.1 ns (mean ± s.e.m., n=8) (**Figure 5A**) and RNA strands bound tightly to ≥100 kDa protein or larger clusters are not expected to rotate appreciably on the 50 ns accessible timescale (26). Segmental motions even within rotationally restricted RNA decrease observed anisotropy to a *B*_0_ value of ∼0.2 within ∼1 ns (**Figure 5A-C**). Thus, the fraction of non-rotating (anisotropic) probes ∼1 ns after the laser pulse, termed *B*/*B*_0_ (see Methods), indicates the proportion of RNA probes bound to helicase or sequestered into the RPCs.

**Figure 5.**
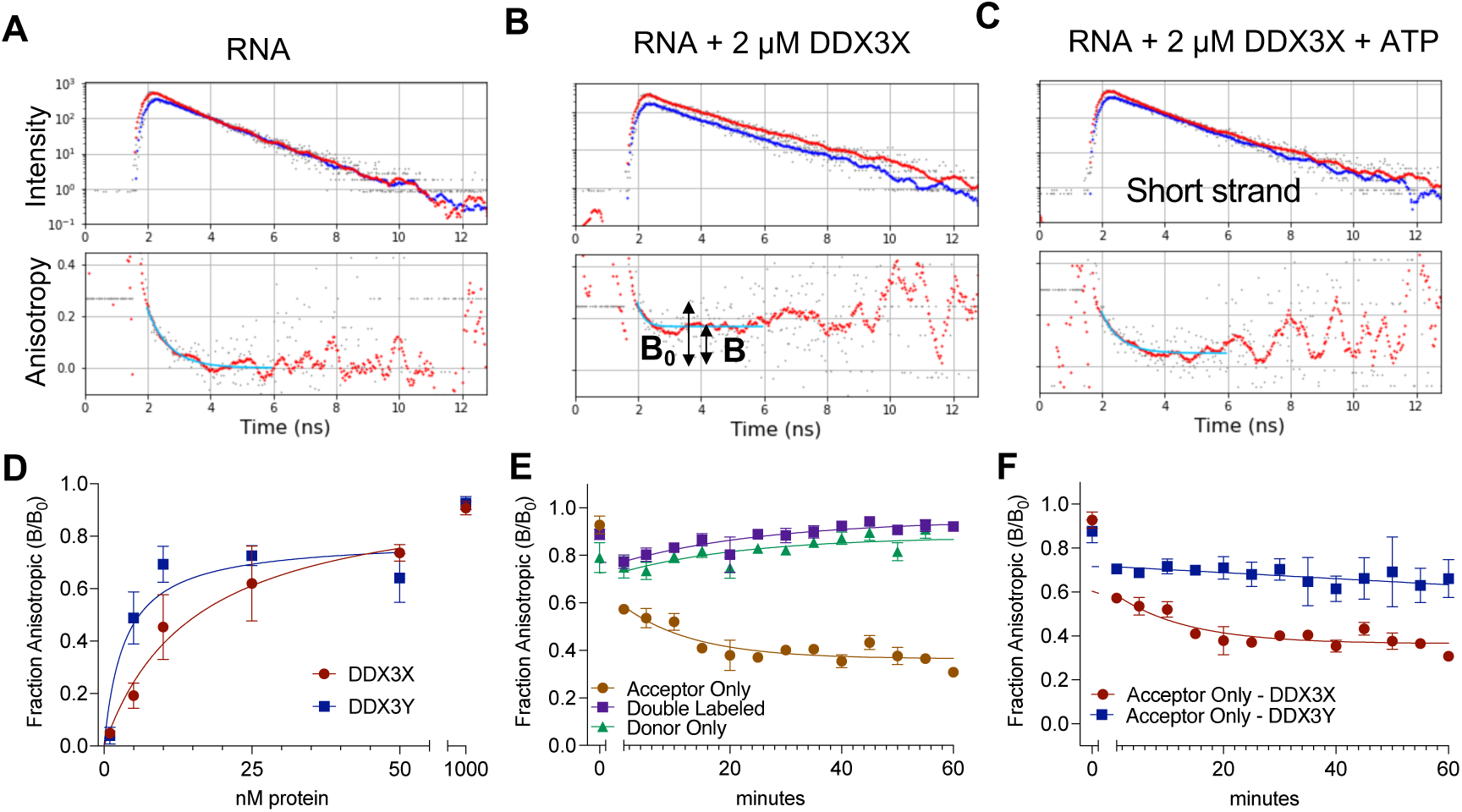
DDX3X has stronger strand release activity than DDX3Y when unwinding double stranded RNA substrate. (A-C) Example time-resolved anisotropy traces of (A) dsRNA alone and (B) dsRNA with 2 µM full-length MBP-DDX3X (short strand shown in graph but plot is representative of all species and (C) and dsRNA with 2 µM full-length MBP-DDX3X and 1 mM ATP (short strand shown in plot). The parallel and perpendicular photon intensities are plotted in the upper panel and the anisotropy decay is plotted in the lower panel. The blue curve is fit to the anisotropy traces with an exponential decay equation. Example traces of the time resolved fluorescence anisotropy of the AO (Alexa647-short strand, acceptor signal) alone, with the addition of DDX3X, and with the addition of ATP to the protein. Lower panel shows the calculated anisotropy which is fit to an exponential decay (blue curve). B_0_ is the segmental motion of the RNA construct calculated from the asymptote B of the RNA with protein recording without ATP addition are shown in the middle panel. (D) Fraction anisotropic (B/B_0_) (slowly tumbling) molecules of RNA with either full-length MBP-DDX3X or DDX3Y at low (nM) concentrations of protein. Values were fit to an exponential decay equation using GraphPad Prism 9. Curves represent three independent measurements for each point. (E) Representative graph showing fraction anisotropic (slowly tumbling) of AO (Alexa647 labeled), DO (Cy3 labeled), and DL (Alexa647 and Cy3 labeled) molecules upon addition of ATP based on time-resolved fluorescence anisotropy analysis. With the addition of ATP, B/B_0_ for the short strand RNA decreases with time and B/B_0_ for DO and DL stays high over time. Values were fit to an exponential decay equation using GraphPad Prism 9. Curves represent three independent measurements for each point. Values were fit to an exponential decay equation using GraphPad Prism 9. Curves represent three independent measurements for each point. (F) Fraction anisotropic (slowly tumbling) of AO (Alexa647 labeled) molecules upon addition of ATP based on time-resolved fluorescence anisotropy analysis for DDX3X and DDX3Y. With the addition of ATP, B/B_0_ for the short strand RNA decreases with time. Values were fit to an exponential decay equation using GraphPad Prism 9. Curves represent three independent measurements for each point. Values were fit to an exponential decay equation using GraphPad Prism 9. Curves represent three independent measurements for each point.

*B*/*B*_0_ is essentially zero in free RNA due to unrestricted tumbling (**Figure 5A**). Within 1 – 3 min after the addition of either DDX3X or DDX3Y, anisotropy of the A647 in the duplex RNA, after the fast segmental rotation, increased to *B*/*B*_0_ = 0.9, indicating that tumbling of most of the molecules became restricted by capture into the RPCs and binding protein (**Figure 5B, 5D, S5A**). Consistent with the diffusion measurements, less DDX3Y than DDX3X was needed to constrain RNA tumbling (**Figure 5D**).

Anisotropy decay also enabled us to determine which short or long product strand is preferentially released after unwinding. The product release step has not been previously assessed for a DEAD-box helicase reaction. *B*/*B*_0_ of the shorter (A647) strand began to drop immediately after addition of ATP (**Figure 5E and F**), while *B*/*B*_0_ of the longer (Cy3) strand (**Figure S5B**) and of DL RNA molecules (**Figure S5C**) remained high (**Figure 5E**). Preference to release the shorter strand was observed for both DDX3X and DDX3Y but the decrease in anisotropy for the shorter AO strand was faster for DDX3X than for DDX3Y (**Figure 5F**). Together, these findings suggest that product strand release, which is slower for DDX3Y than for DDX3X, is a major factor contributing to DDX3X’s faster catalytic cycle relative to DDX3Y’s (**Figure 5**).

### Disease-related mutants of DDX3X reveal a relationship between cluster size and enzyme activity

Lastly, we investigated if RPC size and enzyme activity are correlated. To do so, we utilized nine disease-related point mutants of DDX3X which have variable *in vitro* catalytic activity relative to WT DDX3X (19, 22, 23, 27, 28). We reasoned that these mutants could serve as an experimental substrate to expand the sample size of DDX3 RPCs. To this end, we purified the nine mutants indicated in **Figure S6A** and assessed their diffusion rates (**Figure S6B**) and activity for separating the double-labeled dsRNA I (**Figures S6C and S6D**). When comparing diffusion rate (which decreases with increasing cluster size) to strand separation capacity (separation of double stranded RNA according to decrease in FRET efficiency and decrease in concentration of doubly-labeled particles), we found that the relationship between these parameters displays a maximum at intermediate cluster size. WT DDX3X lies near the peak of the curve (**Figure 6A**). Interestingly, DDX3Y is positioned near the majority of the DDX3X mutants on this curve with slow diffusion (excessive cluster size) and low activity, suggesting that DDX3Y shares these characteristics with the disease-related mutant DDX3X proteins (**Figure 6A**). Additionally, there was a decrease in activity with the increase of diffusion rate past the optimum which tracks well with smaller clusters and decreased activity observed with the double truncation (ΔΔ) constructs. Two mutants (G302V and L505V) were outliers in our analysis of cluster size and strand separation activity. This may indicate that cluster size and strand separation are “uncoupled” in these variants. Characterization of additional constructs that fall outside of this curve might help to identify the factors that lead to this deviation. The cluster size of WT DDX3X RPCs is seemingly optimized for maximum activity (**Figure 6B**), perhaps to enable multiple partial cycles of helicase and re-annealing thus increasing the stochastic probability of full strand separation.

**Figure 6.**
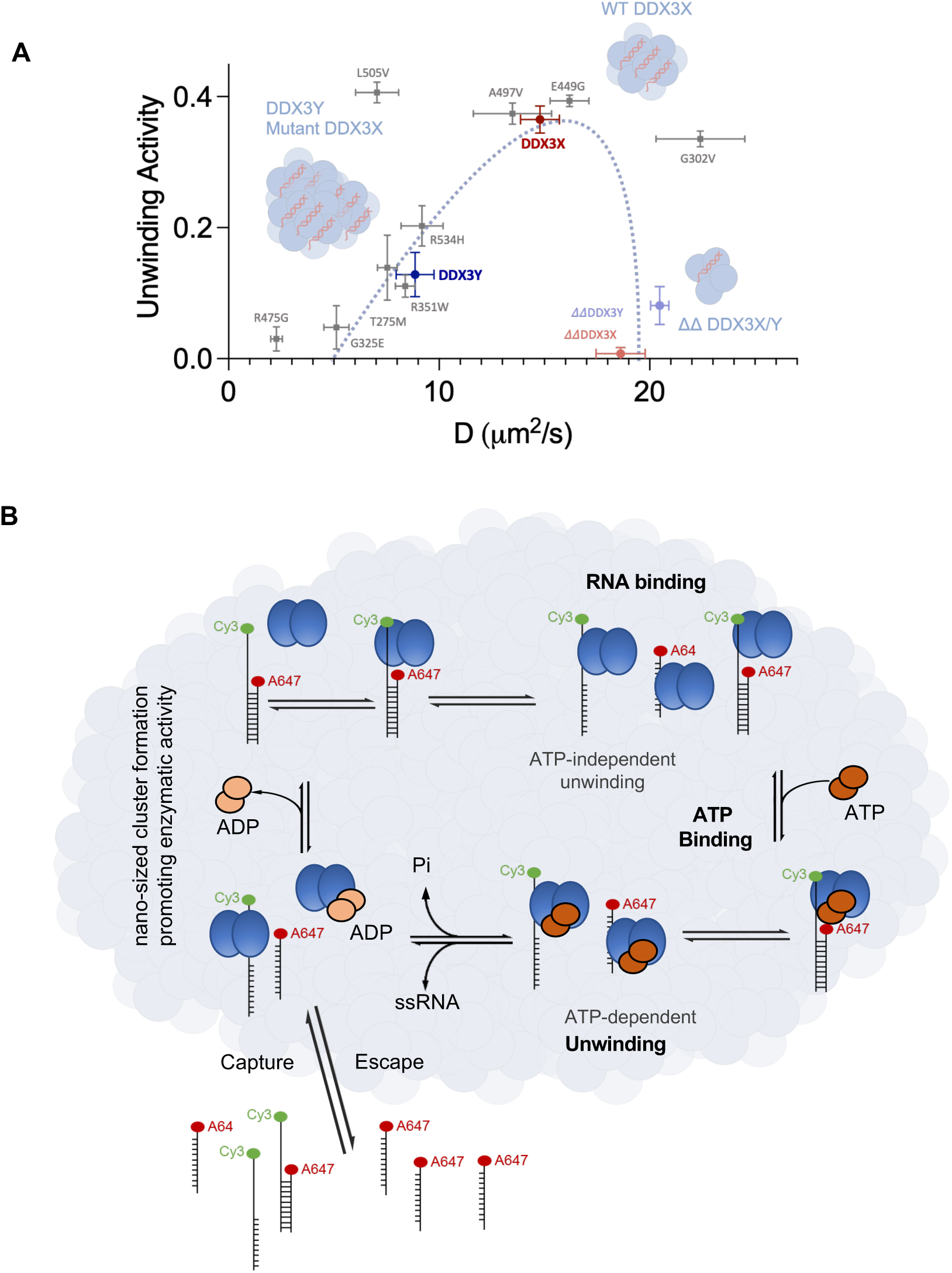
WT DDX3X forms RPCs of the optimal size to maximize enzyme activity, allowing RPCs to act as DDX3 “helicase hubs.” (A) 2D plot of dsRNA I unwinding activity vs. cluster diffusion rates (measured by dcFCCS) for several mutants of DDX3X, WT DDX3X, DDX3Y, and the N- and C-terminal truncation variants of DDX3X and DDX3Y. Unwinding activity was calculated using equation 13 (see Methods) and includes the decrease in double-labeled bursts and decrease in proportion of high FRET DL particles. Overlaid is a schematic line showing the link between catalytic activity and cluster size to guide the eye. Very large and small clusters have low RNA unwinding activity, however midsized clusters (including WT DDX3X) have optimal RNA unwinding activity. (B) Proposed schematic of the enzymatic cycle of DDX3X and DDX3Y and the role of DDX3 RNA-protein clusters. First, RPCs of DDX3X or DDX3Y “capture” free RNA molecules. Once captured, RNA molecules are bound by DDX3 within the cluster (as measured by FRET efficiency increase, time-resolved anisotropy, and EMSAs). Once bound, DDX3 exhibits a low degree of ATP-independent strand separation activity. Upon the addition of ATP, more robust strand separation occurs (as measured by FRET efficiency decrease, decrease in the fraction anisotropic of short strand obtained from time-resolved anisotropy and the gel-based unwinding assays). After ATP hydrolysis, DDX3 releases its product single-stranded RNAs, with a preference for releasing the shorter of the two duplex strands. The formation of RPCs likely increases the local concentration of DDX3 and substrate RNA to facilitate more rapid turnover, as DDX3 is non-processive and must fully repeatedly release its substrate to begin another round of strand separation.

## Discussion

We report the first evidence that the sexually dimorphic RNA helicases DDX3X and DDX3Y form nano-sized clusters, RPCs, both *in vitro* and in cells. The formation of these clusters is driven by the proteins’ intrinsically disordered regions, which also determine the differences in catalytic activity between these homologs. During the catalytic cycle, these differences manifest most strongly at the RNA release step (**Figure 5**), as DDX3X releases its product strand more quickly than DDX3Y.

RPCs form at protein concentrations as low as 5 nM (**Figure 1**), far below the previously established threshold values of 3 – 5 µM for DDX3X/3Y LLPS (6), indicating that they are distinct assemblies from phase separation-mediated RNP granules. This raises a question as to what functions the RPCs may perform. DDX3X is a crucial factor for translation initiation, unwinding secondary structures in 5’-UTRs (29–33), and DDX3Y also has unwinding activities, although weaker than DDX3X (6). Because DDX3X is a non-processive helicase, RNA structures that are longer than ∼13 – 19 bp require at least two separate catalytic cycles to be unwound, in which DDX3X will fully release the RNA and then rebind (34). We suggest a model in that RPCs act as “helicase hubs,” increasing the concentration of DDX3 near substrate mRNAs (**Figure 6**). This may allow the ribosome to scan the 5’-UTR faster during translation initiation, as DDX3 molecules in the cluster are nearby at high concentration to continue unwinding after the initial cycle.

Beyond translation initiation, it has been proposed that DDX3X also “scouts” for endogenous dsRNA species in the cytoplasm, where it unwinds them to avoid a dsRNA-mediated autoimmune response (35). Based on our findings herein, it is possible that DDX3 RPCs perform this scouting function. DDX3 RPC helicase hubs may be analogous to the recently described “transcription hubs,” which assemble the factors necessary for RNA transcription using interactions mediated by intrinsically disordered regions (36). Thus, IDR-enabled RPCs may be the relevant active form of DDX3X and DDX3Y involved in translation initiation, cytoplasmic scouting, and other functions. While *in vitro* RPCs only consist of DDX3 and RNA, cellular RPCs likely contain DDX3, RNA, and partner proteins that aid in the various functions of DDX3X, such as translation factors or ADAR1 (in the case of cytoplasmic scouting (35)). The composition of these RPCs beyond DDX3 merits future study.

The results support a strong correlation between the formation of the RNA-protein clusters and the enhanced helicase activity of the DDX3s. Removing both IDRs from either DDX3X or DDX3Y (ΔΔ constructs) disrupted the RNA unwinding, ATP hydrolysis, RNA binding, and cluster formation of both proteins (**Figures 2 and 3**), suggesting a connection between the formation of the RPCs and catalytic activity. When truncating individual IDRs, we found that the N-terminal IDR differentiated the catalytic activities of DDX3X and DDX3Y, while the C-terminal was equally critical for ATP hydrolysis and RNA unwinding in both proteins. These features align with sequence conservation: the N-terminal IDRs are the most divergent regions of DDX3X and DDX3Y, while the C-terminal IDRs are more similar. When cluster formation was measured for each truncation, we found that the N-terminal IDRs of DDX3X and DDX3Y were driving cluster formation. The C-terminal IDRs drive the difference in the extent of cluster formation, possibly through inter- or intra-protein rearrangement since truncating the C-terminal IDRs does not seem to alter RNA binding affinity.

When we applied our multiparameter confocal technique to several disease-related point mutants of DDX3X, we further revealed the relationship between cluster size and enzymatic activity, as many of the DDX3X mutants show low activity and very large cluster size. The activity and clustering of DDX3Y is comparable to these mutants, suggesting that the combination of its amino acid differences (compared to WT DDX3X) results in characteristics similar to potentially deleterious disease-related DDX3X point mutants (**Figure 6**). A similar phenomenon was recently observed with another pair of sexually dimorphic proteins, the synapse receptors NLGN4X and NLGN4Y (37). While NLGN4X undergoes complete maturation (processing through the endoplasmic reticulum and transport to the synaptic surface), NLGN4Y is maturation deficient, as it is largely retained in the endoplasmic reticulum in a similar manner to autism spectrum disorder-related mutants of NLGN4X. While WT NLGN4X could partially rescue the maturation of mutant NLGN4X when coexpressed, NLGN4Y was unable to do so (37). We propose a similar mechanism may exist for DDX3X and DDX3Y. Coexpression of WT DDX3X (with its optimal clustering and higher activity) with a mutant (as in XX individuals) may lead to less severe phenotypes than coexpression of mutant DDX3X with the mutant-like DDX3Y (as in XY individuals). These findings, together with our concurrent, in-depth study of these disease-related mutants (38), point towards a model in which co-expression of mutant DDX3X variants with DDX3Y exacerbates the deleterious properties of DDX3X mutants (higher than optimal clustering and lower than optimal activity), which may contribute toward sex biases observed in DDX3-related disorders.

The findings of this work were made possible by combining classical ensemble biochemistry techniques with our novel multiparameter confocal spectroscopy approach. Using this powerful technique, we extracted information regarding the diffusion, RNA strand stoichiometry, and anisotropy of particles in addition to detailed information regarding the catalytic cycle of these enzymes, all from the same experiment. This approach revealed that the extent of strand release is a major differentiating step between the catalytic cycles of DDX3X and DDX3Y, with DDX3X releasing its product strand faster than DDX3Y (**Figure 5F**). The difference in strand release likely stems from the difference in ATP hydrolysis speeds between DDX3X and DDX3Y, as DDX3X reaches a faster maximum rate of phosphate release from ATP hydrolysis than DDX3Y (6) and ATP hydrolysis specifically governs strand release after unwinding (39). The ability of this technique to generate such detailed and varied data all at once will undoubtedly prove of great benefit to the study of other DEAD-box helicases and other RNA-processing enzymes beyond the DEAD-box family.

## Acknowledgements

We thank Dr. Yulia Gonskikh for her assistance with the gel-based unwinding assay, Samantha Sustek for her assistance obtaining negative stain TEM images, and the UPenn Electron Microscopy Resource Laboratory for the use of the Tecnai 12 TEM. Schematic figures were created in Biorender. **Funding:** This work was supported by the National Institutes of Health (R35GM133721 and R01HL160726 to K.F.L.; R35GM133721-03S1 to A.Y.; T32GM132039 to A.Y. and M.C.O.; and R35GM118139 to Y.E.G.). K.F.L. is supported by the American Cancer Society (RSG-22-064-01-RMC), the Damon Runyon Innovator Award, and the Linda Pechenik Montague Investigator Award.

## Author contributions

A.Y. and M.C.O. performed the molecular cloning and protein purifications. A.Y. and H.S. performed all-time-resolved spectroscopy, dcFCCS, smFRET, FCS, and fluorescence anisotropy experiments and analyzed the data together with Y.E.G. A.Y. performed mass photometry and analyzed the data together with H.S. and Y.E.G. M.C.O. performed the gel-based unwinding, EMSA and malachite green assays and analyzed the data. M.C.O. assembled samples for negative stain TEM and collected the images. All authors participated in the conception of ideas, planning of experiments, discussion of results, and writing/editing the manuscript.

## Data and material availability

Data are available in the manuscript or the supplementary material. Materials are available upon request to corresponding authors.

## Figures and Figure Legends

**Figure S1.**
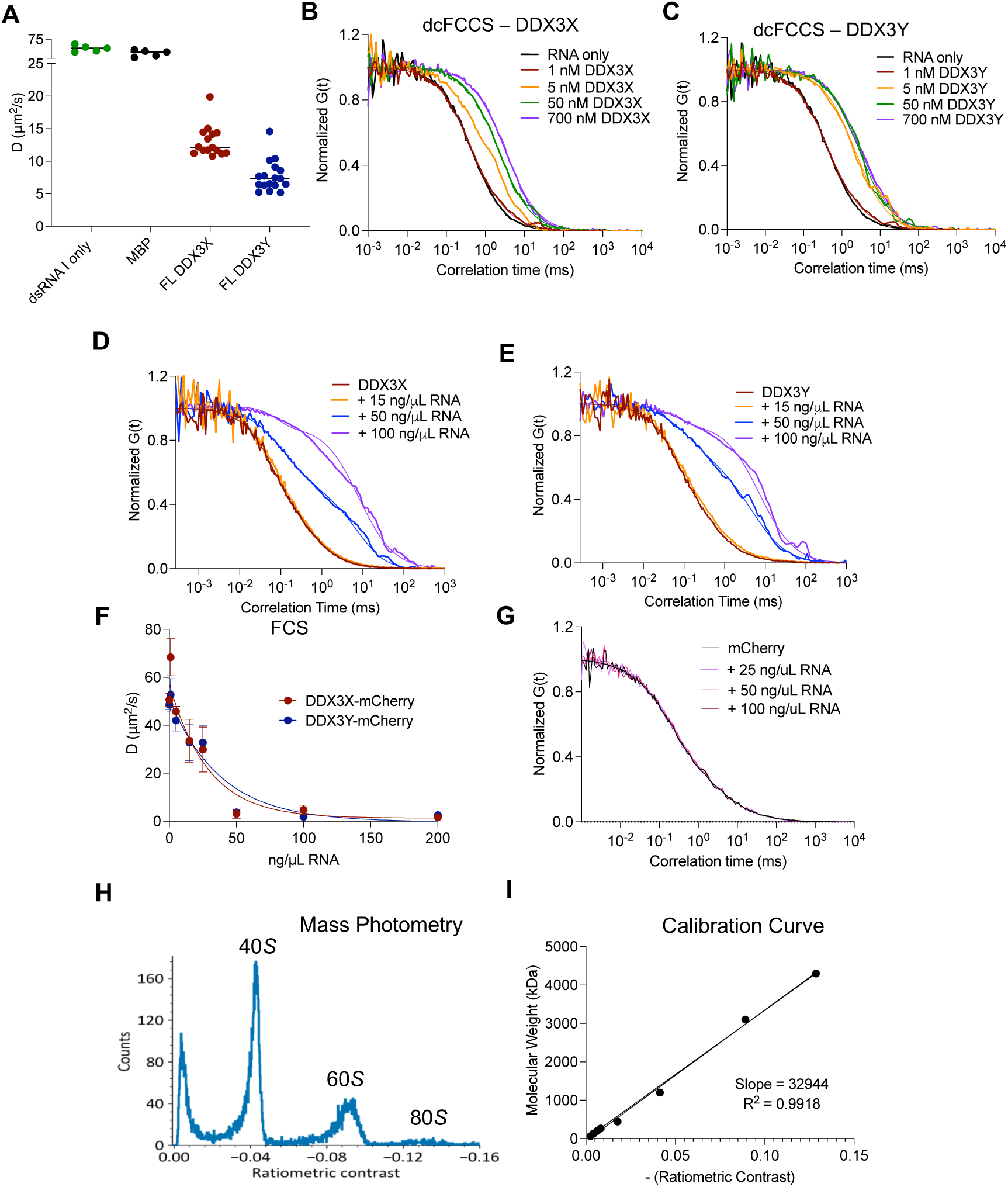
DDX3X and DDX3Y form nano-sized RNA-protein clusters *in vitro,* even at nanomolar concentrations. (A) Diffusion coefficients (µm^2^/s) obtained by fitting the cross-correlation curves using single component diffusion model for RNA alone, MBP protein, full-length MBP-DDX3X, full-length MBP-DDX3Y. These diffusion times correlate with the size of the molecules. (B) Normalized cross-correlation curves showing slower diffusion with increasing concentration of MBP-DDX3X. (C) Normalized cross-correlation curves showing slower diffusion with increasing concentration of MBP-DDX3Y. (D) Auto-correlation curves of mCherry tagged DDX3X at 80 nM of protein with increasing total HEK293T RNA concentrations. The fits deviated from the single diffusing component with increasing HEK293T concentrations. (E) Auto-correlation curves of mCherry tagged DDX3Y at 80 nM of protein with increasing total HEK293T RNA concentrations. The fits deviated from the single diffusing component with increasing HEK293T concentrations. (F) Diffusion coefficients (µm^2^/s) of mCherry tagged DDX3X and DDX3Y with increasing concentrations of HEK293T RNA added. Values were fit to an exponential decay equation using GraphPad Prism 9. Curves represent three independent measurements for each point, error bars represent SEM. (G) Auto-correlation curves of mCherry alone at 80 nM of protein with increasing total HEK293T RNA concentrations. mCherry doesn’t show any evidence of protein binding. (H) Histogram of the ratiometric contrasts of the 80 S ribosome used as a calibrant to estimate the cluster sizes in mass photometry. (I) Molecular weight vs. ratiometric contrast of proteins used for the mass calibration. Values were fit to a linear regression equation using GraphPad Prism 9.

**Figure S2.**
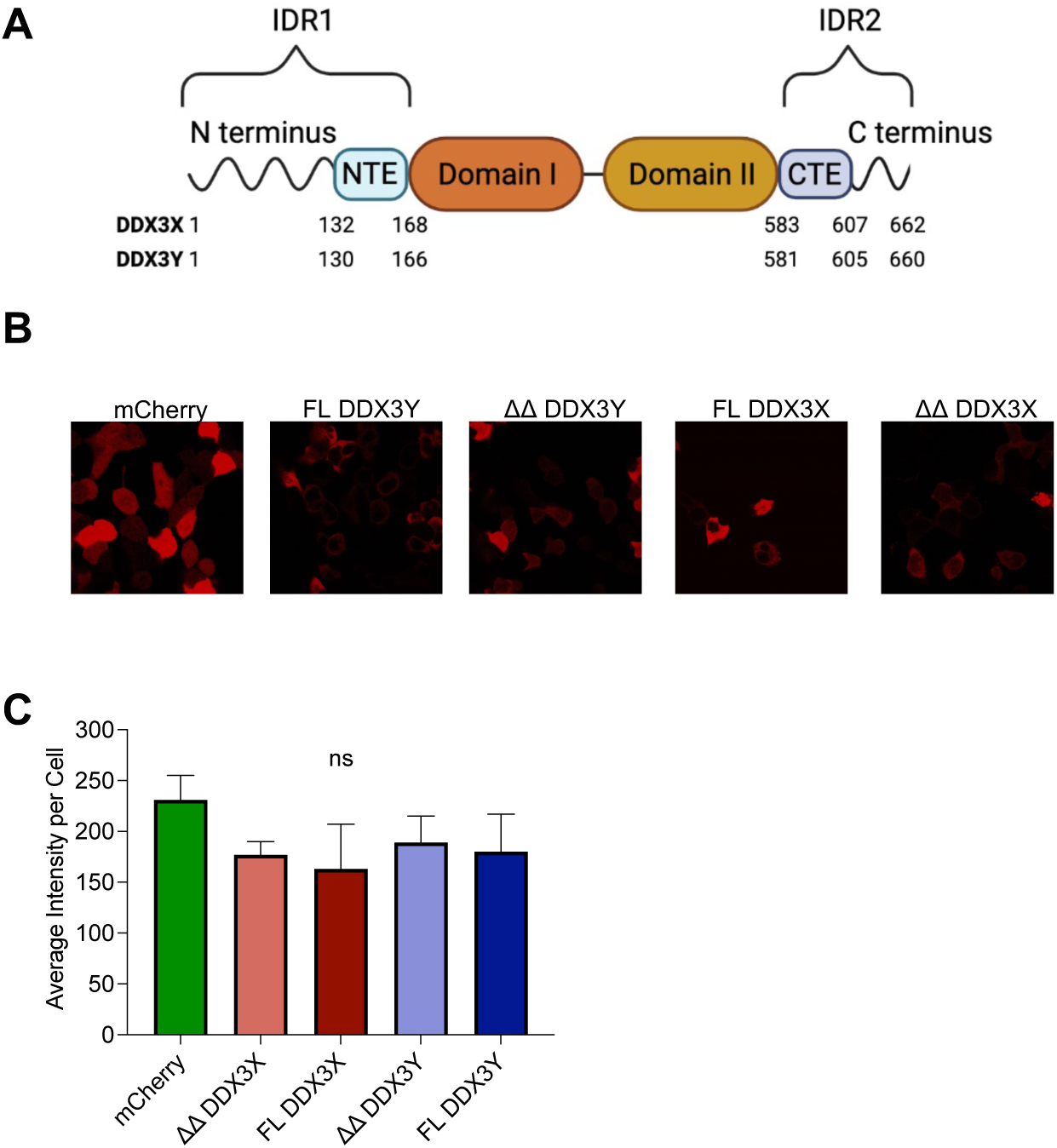
DDX3X and DDX3Y form nano-sized RNA-protein clusters *in vitro*. (A) Schematic of the domain architecture of full-length DDX3X and DDX3Y. (B) Representative images of the total HEK293T cell intensities imaged with a Zeiss Widefield Microscope. (C) Average intensity per cell of mCherry, ΔΔ DDX3X-mCherry, full-length DDX3X-mCherry, ΔΔ DDX3Y-mCherry, full-length DDX3Y-mCherry from live transfected HEK293T cells. Plots and significance were calculated in GraphPad Prism 9 using the Student’s two-tailed t-test. ns, p < 0.12, *, p < 0.033, **, p < 0.002. ***, p < 0.0002. ****, p < 0.0001. (D) Average number of molecules diffusing in the beam in live transfected HEK cells; mCherry, ΔΔ DDX3X-mCherry, full-length DDX3X mCherry, ΔΔ DDX3Y-mCherry, full-length DDX3Y-mCherry. Plots and significance were calculated in GraphPad Prism 9 using the Student’s two-tailed t-test. ns, p < 0.12, *, p < 0.033, **, p < 0.002. ***, p < 0.0002. ****, p < 0.0001.

**Figure S3.**
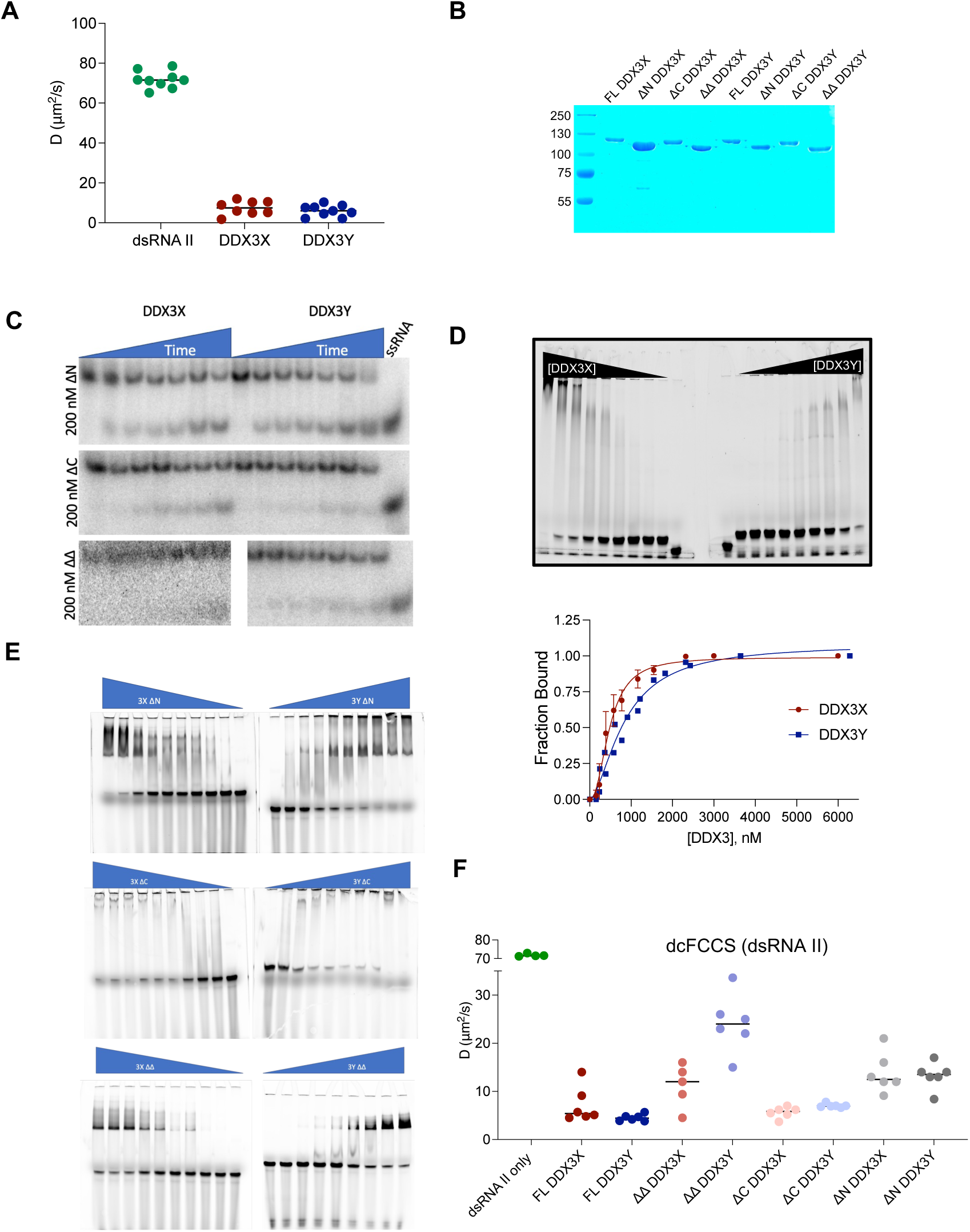
Elucidating the role of IDRs in the catalytic activity of DDX3X and DDX3Y *in vitro*. (A) Diffusion coefficients (µm^2^/s) obtained by fitting the cross-correlation curves using single component diffusion model for dsRNA II alone, and dsRNA II with full-length MBP-DDX3X, full-length MBP-DDX3Y at 200 nM. These diffusion times correlate with the size of the molecules. (B) Coomassie-stained SDS-PAGE gel showing purity of each MBP-tagged protein. (C) Representative unwinding gel for the indicated constructs at 200 nM concentration. (D) Upper, representative EMSA of full-length MBP-DDX3X/DDX3Y with varying protein concentrations, showing binding of the protein to the RNA protein using the fluorescent duplex RNA. Lower, quantification of EMSAs for full-length MBP-tagged proteins. Values were fit to Hill kinetics using GraphPad Prism 9. Curves represent three independent measurements for each point. (E) Representative EMSA gel of each truncated construct. (F) Diffusion coefficients (µm^2^/s) obtained by fitting the cross-correlation curves using single component diffusion model for dsRNA II alone, and dsRNA II with MBP-tagged full-length DDX3X, full-length DDX3Y, ΔΔ DDX3X, ΔΔ DDX3Y, ΔC DDX3X, ΔC DDX3Y, ΔN DDX3X, and ΔN DDX3Y at 2 µM. These diffusion coefficients correlate with the size of the molecules.

**Figure S4.**
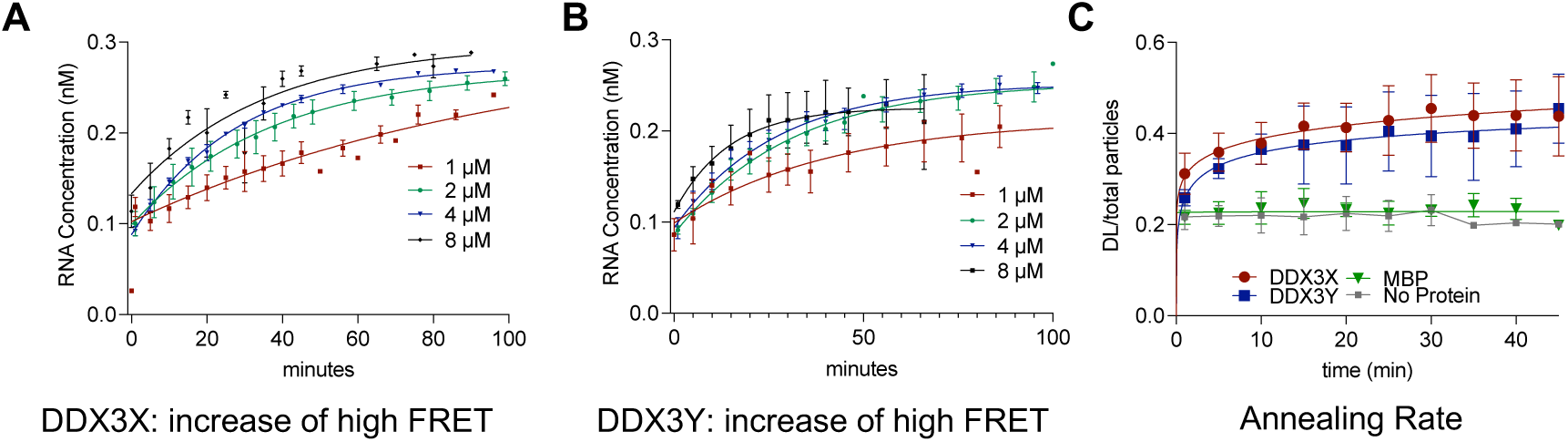
DDX3X and DDX3Y have similar RNA reannealing activities when measured by a single molecule approach. (A) Individual kinetics curves plotted for 1, 2, 4, 8 µM full-length MBP-DDX3X, accounting for the proportion of DL bursts, the proportion of high FRET bursts, and starting concentration of RNA (0.5 nM, see methods, equations 5 and 6). RNA concentration is plotted on the Y-axis. Values were fit to an exponential decay equation using GraphPad Prism 9. Curves represent three independent measurements for each point. (B) Individual kinetics curves plotted for 1, 2, 4, 8 µM full-length MBP-DDX3Y, accounting for the proportion of DL bursts, the proportion of high FRET bursts, and starting concentration of RNA (0.5 nM, see methods, equations 5 and 6). RNA concentration is plotted on the Y-axis. Values were fit to an exponential decay equation using GraphPad Prism 9. Curves represent three independent measurements for each point. (C) Annealing experiment showing how both single-stranded RNAs (either Alexa647 or Cy3 labeled) were combined and the proportion of DL RNA was monitored either with no protein added or with the addition of 4 µM MBP, MBP-tagged 4 µM full-length DDX3X, or 4 µM full-length DDX3Y over the course of 60 minutes. Values were fit to an exponential decay equation using GraphPad Prism 9.

**Figure S5.**
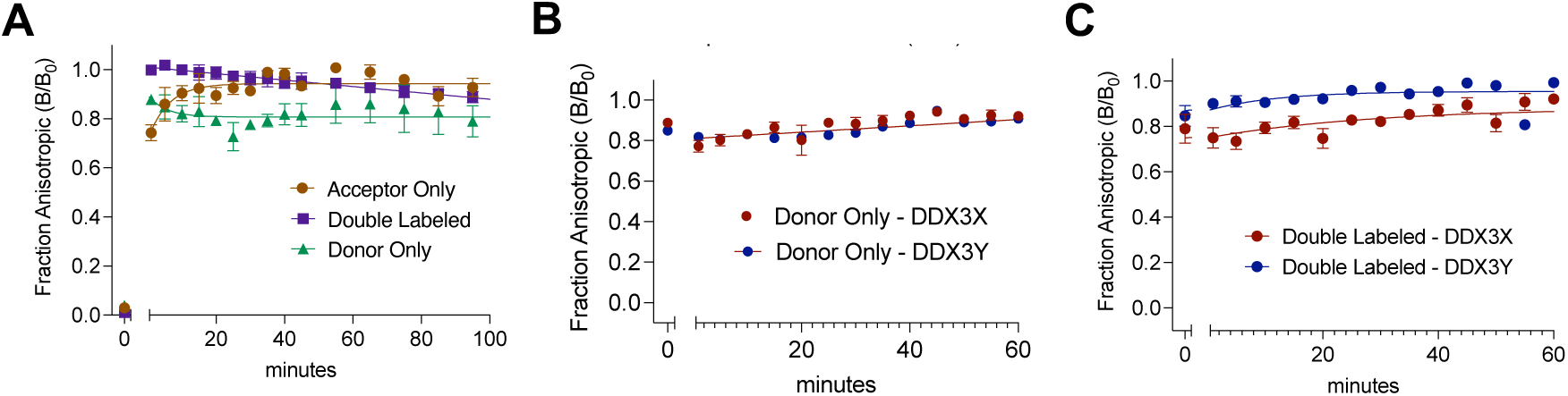
DDX3X and DDX3Y stay bound to double stranded and long strand RNA throughout the catalytic cycle. (A) Fraction anisotropic (slowly tumbling) of AO (Alexa647 labeled), DO (Cy3 labeled), and DL (Alexa647 and Cy3 labeled) molecules for protein only based on time-resolved fluorescence anisotropy analysis. For all species, B/B_0_ stays high over time. Values were fit to an exponential decay equation using GraphPad Prism 9. Curves represent three independent measurements for each point. (B) Fraction anisotropic (slowly tumbling) of DO (Cy3 labeled) molecules upon addition of ATP based on time-resolved fluorescence anisotropy analysis. Long strand RNA has high B/B_0_ when protein and ATP are added. Values were fit to an exponential decay equation using GraphPad Prism 9. Curves represent three independent measurements for each point. Curves represent three independent measurements for each point. _(C)_ Fraction anisotropic (slowly tumbling) DL (Cy3 and Alexa647 labeled) molecules upon addition of ATP based on time-resolved fluorescence anisotropy analysis. Duplex RNA has high B/B_0_ when protein and ATP are added. Values were fit to an exponential decay equation using GraphPad Prism 9. Curves represent three independent measurements for each point.

**Figure S6.**
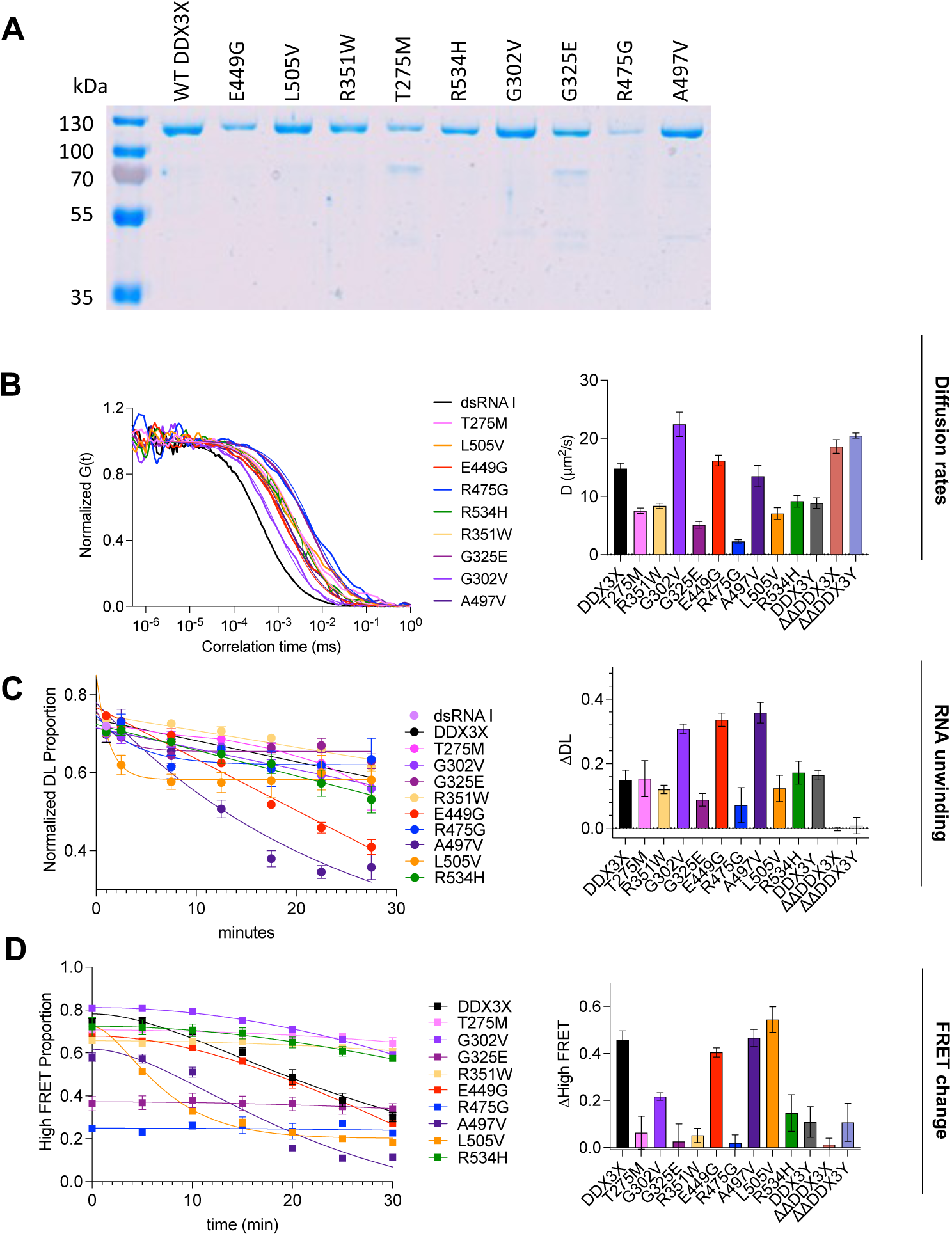
DDX3X mutants cluster size is correlated with their ability to unwind dsRNA. (A) Coomassie-stained SDS-PAGE gel showing purity of each MBP-tagged DDX3X mutant protein along with WT DDX3X. (B) Normalized cross-correlation curves of fluorescent dsRNA I diffusing with the indicated DDX3X mutant (left). Diffusion coefficients (µm^2^/s) obtained by fitting the cross-correlation curves using single component diffusion model for the indicated mutant (right). (C) Plot showing the proportion of DL (duplex, both Alexa647 and Cy3) bursts with the addition of ATP (left panel). Values were obtained by quantifying the respective burst proportion (see method). Curves represent three independent measurements for each point (left). ΔDL is shown on the right panel and was calculated using equation 11 (see method). (D) Plot showing the proportion of high FRET with the addition of ATP. Curves represent three independent measurements for each point (left panel). ΔHigh FRET is calculated using equation 12 (see methods) shown on the right panel. Each bar represents three independent measurements.

