## Supplemental Text for "DDX3X and DDX3Y constitutively form nano-sized RNA-protein clusters that foster enzymatic activity"

### Methods

**Protein expression and purification**: MBP-tagged and mCherry-tagged protein expression vectors were generated as previously described (6). All plasmids were transformed into BL21-CodonPlus(DE3)-RIL cells (Agilent) for protein expression. Cells were then grown to OD 0.6 at 37° C while shaking at 180 rpm. Once OD 0.6 was reached, protein expression was induced by the addition of 1 mM IPTG. Induced cultures were then grown at 16°C for 16 hours while shaking at 180 rpm. After 16 hours, cultures were harvested by centrifugation and resuspended in a lysis buffer consisting of 25 mM Tris-HCl pH 7.5, 500 mM NaCl and 1 mM PMSF. Resuspended cells were then lysed by sonication at 4° C. Following sonication, lysates were clarified by centrifugation and the applied to a Ni-NTA resin column for immobilized metal affinity chromatography purification. Once bound, the resin was washed with lysis buffer supplemented with 50 mM imidazole, then by 25 mM Tris-HCl pH 7.5 and 2 M NaCl. After washing, protein was eluted from the resin using lysis buffer supplemented with 500 mM imidazole. Eluate was concentrated and was purified via size exclusion chromatography using a Superdex 75 resin column on an Äkta pure HPLC (Cytiva). Sizing for MBP proteins was carried out at 4° C, while sizing for mCherry proteins was carried out at room temperature. Proteins were eluted from the sizing column using protein storage buffer (25 mM Tris-HCl pH 8.0, 200 mM NaCl for MBP-tagged proteins and 25 mM Tris-HCl pH 8.0, 500 mM NaCl, 10% glycerol, 2 mM DTT for mCherry-tagged proteins). Eluates were then concentrated, aliquoted, flash frozen in liquid nitrogen, and stored at -80° C until use. Protein purity was examined using SDS-PAGE and by spectrophotometry to measure $A_{260}/A_{280}$ ratio to indicate RNA carry over. MBP-tagged proteins had a $A_{260}/A_{280}$ of ~ 0.75, while mCherry-tagged proteins had a ratio of ~ 0.55.

**Gel-based RNA unwinding assay:** RNA unwinding assays were performed as previously described (21). Briefly, a 13-mer RNA strand (5'-AGCACCGUAAAGC-3') (Eurofins Genomics) was 5' radiolabeled using T4 polynucleotide kinase (PNK). After purification of labeled product, labeled 13-mer RNA was annealed to a 38-mer (5'-GCUUUACGGUGCUUAAAACAAAACAAAACAAAACAAAA-3') (Eurofins Genomics) in 1X duplex annealing buffer (10 mM MOPS pH 6.5, 1 mM EDTA, 50 mM KCl) to produce RNA II. Trace amounts of this radiolabeled duplex were incubated for 5 minutes at room temperature with the indicated concentration of MBP-DDX3X or MBP-DDX3Y in 1X helicase reaction buffer (40 mM Tris-HCl pH 8.0, 50 mM NaCl, 2.5 mM $MgCl_2$, 0.01% NP-40 alternative, 2 mM DTT, and 0.1 U/mL RNase inhibitor) in a 30 µL reaction. After this preincubation, 3 µL of the reaction was removed and quenched with 3 µL of 2X helicase reaction stop buffer (50 mM EDTA, 1% SDS, 20% glycerol, and trace amounts of bromophenol blue and xylene cyanol) and stored on ice. To initiate the reaction, 3 µL of 20 mM ATP was added (2 mM final concentration). The reaction was mixed thoroughly and placed at 37°C. 3 µL were removed from the reaction at 15 s, 30 s, 1 min, 2 min, 3 min, and 6 min, mixed with 3 µL of 2X helicase reaction stop buffer and placed on ice. After 6 minutes, 3 additional µL were removed from the reaction, mixed with stop buffer, and heated to 95°C for two minutes to serve as a single stranded marker. Samples were then loaded into a 20% 19:1 acrylamide gel made in 0.5X TBE buffer and run at 140 V for 1:40 hrs in 1X TBE run buffer. Gels were dried at 80°C for one hour under vacuum, then exposed to a phosphor screen for 36 hours. Images were taken using an Amersham Typhoon using the phosphorimaging settings. Band intensities were measured using Fiji. For each concentration, the fraction unwound was calculated using Equation 1:

$$Fraction\ unwound = 100 * (\frac{I_{ss}}{I_{ss} + I_{ds}}) \quad [1]$$

Where, $I_{ss}$ and $I_{ds}$ are the intensities of the single-stranded and double-stranded bands at each time point. Fraction unwound values were plotted versus time in GraphPad Prism 9 and fit with a single exponential rate equation to calculate the $K_{unw}$ for each concentration. $K_{unw}$ were then plotted versus protein concentration and fit with the Hill equation to get the $K_{unw}^{max}$ and the $K_{1/2}$.

**Electrophoretic mobility shift assays (EMSAs) for protein/RNA binding**: EMSAs were performed by mixing the indicated concentration of MBP-DDX3X, MBP-DDX3Y, or truncated construct with 100 nM duplex RNA I in a buffer of 50 mM Tris-HCl pH 7.5, 150 mM KCl, 2 mM $MgCl_2$, 100 mM beta-mercaptoethanol and 0.1 mg/mL BSA. Reactions were mixed and left at room temperature for 30 minutes. 1 µL of 5X Novex Hi-Density sample buffer (ThermoFisher) was added and the reactions were run on a 6% 37.5:1 acrylamide made in 0.5X TBE gel for 1:50 hrs at 120 V in 0.5X TBE run buffer. Gels were then imaged using an Amersham Typhoon on the "Cy5" channel. Band intensities were measured using Fiji, and fraction bound was calculated using Equation 2:

$$Fraction\ bound = 100 * (1 - (\frac{I_n}{I_o})) \qquad [2]$$

Where, $I_n$ is the intensity of the free probe band at a given concentration and $I_o$ is the intensity of the free probe band at 0 nM protein. Fraction bound was plotted versus concentration in GraphPad Prism 9 and fit with the Hill equation to give $K_D$.

**Malachite green ATPase assays**

To measure the ATPase activity of DDX3X, DDX3Y, and all truncations, a malachite green colorimetric phosphate assay was used as previously described (6) (Sigma-Aldrich). Briefly, proteins were diluted to 1 µM in assay buffer (25 mM Tris-HCl pH 8.0, 200 mM NaCl, 2 mM $MgCl_2$, 1 mM DTT) and mixed with 100 ng/µL total RNA extracted from HEK293T cells. The protein and RNA were allowed to equilibrate for 15 minutes at room temperature before the addition of 2 mM ATP to start the reaction. Reactions were incubated for 30 minutes at room temperature, then quenched with malachite green. Following a further 30 minutes of room temperature incubation, reactions were loaded into a clear-bottom 384 well plate and absorbance of the malachite green at 660 nm was read on a plate reader (Tocris Bioscience). Absorbance values were converted to phosphate released per DDX3 using a standard curve, and background values (calculated from reactions incubated without RNA) were subtracted from each measurement before plotting in GraphPad Prism 9. Significance was determined using Student's two-tailed t test.

**Multiparameter confocal time resolved spectroscopy**

All single-molecule fluorescence burst measurements were performed by multi-parameter confocal time-correlated single photon counting (TCSPC) microscopy and spectroscopy (MicroTime-200; PicoQuant, GmbH). All single molecule measurements were done in FRET buffer (50 mM Tris pH 7.5, 125 mM NaCl, and 2 mM MgCl2) and utilized RNA I: a 18-mer (5'-biotin/ACCGCUGCCGUCGCUCCG/AlexF647N/-3') annealed to a 42-mer (5'-/Cy3/UUUUUUUUUUUUUUUUUUUUUUUUCGGAGCGACGGCAGCGGU-3') (IDT). The donor (Cy3) and acceptor (Alexa647) dyes on the labelled RNA were excited with 532 nm or 560 nm and 637 nm pulsed diode lasers (LDH-D-TA-532, LDH-D-TA-560, LDH-D-TA-637, PicoQuant) operating at 20 MHz in pulsed-interleaved excitation (PIE) mode, through an excitation dichroic filter ZT532/637 (Chroma Technology) and an Olympus UPLanSApo 60x/1.2 w/ water immersion objective lens. Power of both the lasers was kept below 20 μW and the laser was focused 20 μm above the coverslip interface for the measurement. Fluorescence signals from the sample were separated into vertical and horizontal polarized paths by a polarizing beam splitter after passing through a 50 μm pinhole conjugate to the sample plane. Identical dichroic filters (T635 lpxr, Chroma Technology) were used in each pathway to split signals into Cy3 and Alexa647 emission and further selected by bandpass filters ET582/64 (for donor, Cy3) or ET690/70 (for acceptor, Alexa647). Polarized and wavelength-selected photons were projected onto four single-photon avalanche photodiode (SPAD) detectors and cataloged by a HydraHarp TCSPC time-interval analyzer in (PicoQuant). Data was recorded for the labeled RNA alone and then several 5-minute periods (over the course of 1 -2 hrs) after adding the protein to the RNA followed by further data collection (over the course of 1 hr) after adding 1 mM MgATP. *In vitro* measurements were done using Nunc Lab-Tek chambers (ThermoFisher-155411) with borosilicate coverslip bottoms. These chambers were passivated by treatment with 50% (w/v) PEG-8000 solution, incubated at room temperature for 3 – 4 hours, followed by 3 – 4 washes with 300 µL FRET buffer. Rhodamine6-G, atto565 and atto637N dyes were used as standard samples to estimate the confocal detection volume for the 532 nm, 560 nm and 637 nm lasers.

**Single molecule FRET (smFRET):** The acquired files of the PIE data were converted into the universal photon HDF5 file format using the phconvert module of the FRETBursts software (10). Histograms of the photon arrival times in these photon streams were accumulated in, typically, 5-minute recordings. For the FRET analysis, donor and acceptor counts from the two polarized channels were combined. Background correction was applied in each 30 s measurement window and photon bursts above the background were identified and corrected for cross-channel leakage, differential laser power, probe absorptivity, quantum yield and detector sensitivity (40). In the 0.5 – 1 nM concentrations range of labeled RNA or protein, the mean number of molecules in the detection beam was 0.3 – 0.4. Photon bursts corresponding to individual fluorescent particles diffusing through the beam were identified by the sliding window algorithm of m = 8, threshold rate F = 5 times larger than the background rate and threshold of L = 20 in the FRETBursts package.

The three photon streams for each single molecule event after all the corrections are:

donor emission after donor excitation: $F_{Dem/Dex}$

acceptor emission after donor excitation: $F_{Aem/Dex}$

acceptor emission after acceptor excited: $F_{Aem/Aex}$

The FRET efficiency ($E$) and stoichiometry ($S$) were then calculated for each burst as follows:

$$E = \frac{F_{A_{em}/D_{ex}}}{F_{D_{em}/D_{ex}} + F_{A_{em}/D_{ex}}} \quad [3]$$

$$S = \frac{F_{D_{em}/D_{ex}} + F_{A_{em}/D_{ex}}}{F_{D_{em}/D_{ex}} + F_{A_{em}/D_{ex}} + F_{A_{em}/A_{ex}}} \quad [4]$$

*E* gives an estimate of the distance (*R*) between the two probes and *S* gives the probe stoichiometry of each particle: *S* = 1.0 for donor-only (DO) particles; *S* = 0.5 for doubly-labeled (DL) particles; and *S* = 0.0 for acceptor-only (AO) particles. DO, AO and DL populations are selected using the *E vs. S* representation and burst selection filters in FRETBursts. The three types of particles are represented in the main text figures in terms of their RNA concentrations using Equation 5:

$$R_i = \frac{X_{Bursts\_i}}{Tot_{Bursts}} * dsRNA_{conc} \quad [5]$$

Where, $R_i$ is the RNA concentration of the $i^{th}$ population, $X_{Bursts\_i}$ is the DO or AO or DL population after burst categorization, $Tot_{Bursts}$ is the total number of bursts in the *E vs. S* histogram, given as DO + DL + AO, $dsRNA_{conc}$ is the starting dsRNA substrate concentration (0.5 nM).

The DL particles were further sorted into high FRET (HF) and low FRET (LF) components based on *E* value either above or below *E* = 0.5. The DL population undergoing high FRET is then represented in terms of dsRNA concentrations (nM) using Equation 6:

$$R_{HF} = \frac{X_{HF}}{Tot_{HF+LF}} * R_{DL} \quad [6]$$

Where, $R_{HF}$ is the dsRNA concentration (nM) undergoing high FRET, $X_{HF}$ is the occupancy of the high-FRET group within the FRET distribution, $Tot_{HF+LF}$ is the total occupancy of high and low FRET DL components.

**Dual-color Fluorescence Cross Correlation Spectroscopy (dcFCCS):** In dcFCCS, the fluorescence fluctuations of the two spectrally discrete donor and acceptor detection channels are cross-correlated indicating diffusion together, as in a doubly-labeled dsRNA or larger cluster of fluorescent strands. The donor photons during donor excitation ($F_{DexDem}$) and acceptor photons during acceptor excitation ($F_{AexAem}$) within the selected DL particles were cross-correlated in time. Cross-correlation curves were fitted by a diffusion model equation with one or two diffusing components (Equation 7):

$$G(t) = \sum_{i=1}^{n_{Diff}} \frac{\rho[i]}{\left[1 + \frac{t}{\tau_{Diff}[i]}\right][1 + \frac{t}{\tau_{Diff}[i]k^2}]^{0.5}} \quad [7]$$

Where, $\rho$ is the contribution of the $i^{th}$ diffusing species, $k$ is length to diameter ratio of the focal volume, $\tau_{Diff}$ is the mean diffusion time of the $i^{th}$ diffusing species and $n_{Diff}$ is the number of independently diffusing species.

**Fluorescence Correlation Spectroscopy (FCS):** The MBP- and mCherry-tagged DDX3 samples were excited with a 560 nm pulsed diode laser (LDH-D-TA-560, Picoquant). The fluorescence signals were separated by a polarizing beam splitter and projected onto two SPADs, which were then utilized to calculate the auto-correlation curves. Use of two detectors eliminates the effect of detector after-pulses which are not correlated. Data was recorded for several five minutes periods.

**In cell FCS:** HEK293T were cultured in DMEM+ GlutaMAX (GIBCO) with 10% FBS (GIBCO) and 1% Pen/Strep (Corning) in a humidified incubator with 5% $CO_2$ at 37ºC. Cells were counted with a hemacytometer and 300,000 cells were plated into a 6-well plate on day 1. On day 2, cells were transfected with pPB plasmids expressing DDX3X-mCherry, DDX3Y-mCherry, ΔΔ DDX3X-mCherry, ΔΔ DDX3Y-mCherry, or mCherry alone. For the transfection, 1 µg of plasmid was added to 250 µL OPTIMEM. The solution was vortexed for 15 seconds and centrifuged for 1 minute at 500 x g. Then, 1.4 µL Avalance Omni transfection reagent was added. The solution was vortexed for 15 seconds and centrifuged for 1 minute at 500 x g, the solution incubated for 15 minutes. 50 µL of the solution was added dropwise to the well. After 16 hours, media was removed, cells were washed with PBS and FluoroBrite DMEM medium with 10% FBS was added. The cells were imaged in an environmental chamber which kept the samples at 37ºC, 5% $CO_2$, and added humidity. During the experiment, cells were chosen manually under brightfield observation by moving the microscope x-y stage. Cells with very high fluorescence were avoided. Laser power was adjusted to balance the acquisition of sufficient numbers of fluorescent photons to adequately fit the correlation curves while avoiding photo-bleaching and damage of the cells. The laser beam was focused 2 µm above the coverslip interface for in cell measurements. In control experiments, non-transfected cells produced very little uncorrelated auto-fluorescence. Auto-correlation curves were fitted with a diffusion model containing two diffusing species including the mCherry triplet state (Equation 8):

$$G(t) = \left[1 + T\left[exp\left(-\frac{t}{t_{trip}}\right) - 1\right]\right] \sum_{i=1}^{2} \frac{\rho[i]}{\left[1 + \frac{t}{\tau_{Diff}[i]}\right][1 + \frac{t}{\tau_{Diff}[i]k^2}]^{0.5}} \qquad [8]$$

Where, $i$ identifies the faster or slower diffusing species, $\rho$ is the contribution of the $i^{th}$ diffusing species, *k* is length to diameter ratio of the focal volume, $\tau_{Diff}$ is the diffusion time of the $i^{th}$ diffusing species, $T$ is triplet-state fraction and $t_{trip}$ is triplet relaxation time of the fluorophore. Diffusion time, obtained as correlation time at half-maximum amplitude ($\tau_{1/2}$) of the triplet-corrected auto-correlation curve, is used to calculate the apparent diffusion coefficient ($D_{app}$) using the relation, $D_{app} = w_0^2/4\tau_{1/2}$, where $w_0$ is the detection volume radius at the beam waist.

To adjust the transfection conditions to obtain approximately equal fluorescence intensity among the variant constructs, the overall fluoresce intensity of the cells was measured with a

Zeiss widefield epi-fluorescence microscope. The total fluorescence intensity was quantified in ImageJ and the number of cells in an image was counted. The average intensity per cell was quantified for each condition.

**Time-resolved fluorescence anisotropy:** The parallel and perpendicular ns fluorescence intensity decays for DO particles, DL particles and AO particles within the respective selected regions of *E vs. S* plots and burst selection filters were used to calculate the time-resolved anisotropy decay ($r(t)$) using Equation 9. DO particles used donor excitation ($F_{DexDem}$). DL and AO particles used acceptor excitation ($F_{AexAem}$).

$$r(t) = \frac{I_{II}(t) - G\,I_{\perp}(t)}{I_{II}(t) + 2\,G\,I_{\perp}(t)} \quad [9]$$

Where, $r$ is the anisotropy, $I_{II}(t)$ is the parallel fluorescence intensity, $I_{\perp}(t)$ is the perpendicular fluorescence intensity, $G$ is the correction factor for difference in sensitivity of the two detectors. The time resolved anisotropy curves ($r(t)$) were fitted with Equation 10 to obtain rotational correlation time ($\Phi$) and the proportion molecules with hindered rotation ($B/B_0$):

$$r(t) = (B_0 - B)\,exp^{-\frac{t}{\Phi}} + B \quad [10]$$

Where, $B_0$ is the anisotropy value ~1 ns after the laser pulse and completion internal (segmental) motions within the RNA construct. The value of $B_0$ was determined from the asymptote ($B$) of recordings with RNA and protein before adding ATP.

**Mass photometry:** The mass photometry experiments were carried out on a Two$^{MP}$ mass photometer (Refeyn, Oxford, UK) at room temperature. Homemade chambers assembled using silastic sheet (thickness 3 mm) and microscope coverslips (#1.5, 24x50 mm$^2$, Thermo Fisher) were thoroughly cleaned with ethanol, methanol, and distilled water. The chambers were then dried under clean nitrogen. The focus position of the microscope objective was stabilized and locked using filtered FRET buffer before adding the RNA/protein sample for each measurement.

Standard samples (bovine serum albumin (BSA, 66 KDa), yeast alcohol dehydrogenase (150 KDa), Apoferritin from horse spleen (443 KDa) and 40*S* ribosome subunit, 60*S* ribosome subunit, and 80*S* ribosome (1.65, 2.75, 4.40 MDa, respectively) were used to generate interferometric contrast *vs.* molecular mass calibration curves for each experimental day. All the standard samples were diluted to 10 nM and the data was acquired with a 12 x 17 $\mu m^2$ field of view and the collected for 1 minute at 135.1 Hz frame rate. The exposure time was reduced from a default of 7.7 ms to 5.5 ms to avoid saturating the camera with the scattering contrast signal for larger MDa sized particles. The resulting image data was analyzed using Discover$^{MP}$ software provided by the instrument manufacturer. Histograms of interferometric contrast for each landing event were fit by Gaussian components in Discover and the peaks of the Gaussian curves for each species were converted to apparent molecular mass using the calibration curve measured on that experimental day. Relative numbers of each type of particle were quantified by integrating its Gaussian component.

**Negative stain electron microscopy**: 50 µL samples were generated by diluting the indicated construct to 1 µM with 1 nM RNA I in 1X FRET buffer. 3 µL of each sample were then applied to a glow-discharged carbon-coated 300-mesh copper grid (Electron Microscopy Sciences). The gird was stained for 1 minute with 2% uranyl acetate, blotted, stained again, and blotted again. Samples were imaged at room temperature on an FEI Tecnai 12 transmission electron microscope at an acceleration voltage of 100 kV.

**DDX3X Mutants - Multiparameter confocal time resolved spectroscopy and smFRET:**
Change in concentration of double-labeled diffusing particles, $\Delta DL$, was calculated with the following equation:

$$\Delta DL = DL_{t=30\,min} - DL_{t=0\,min} \qquad [11]$$

Where, $DL_{t=30}$ is the concentration of DL bursts quantified 30 minutes after the addition of ATP and $DL_{t=0}$ is the DL bursts before the addition of ATP.

ΔHigh FRET ($\Delta HF$) was calculated with the following equation:

$$\Delta HF = HF_{t=30\,min} - HF_{t=0\,min} \quad [12]$$

Where, $HF_{t=30}$ is the proportion of DL particles with FRET Efficiency over 0.5 quantified 30 minutes after the addition of ATP and $HF_{t=0}$ is the proportion of DL particles with FRET Efficiency over 0.5 before the addition of ATP.

"Unwinding activity" was calculated using the following equation:

$$Unwinding\ Activity = (HF_{t=30\,min} \times DL_{t=30\,min}) - (HF_{t=30\,min} \times DL_{t=30\,min}) \quad [13]$$
